# The effect of Vitamin D (1,25-(OH)_2_-D3) on human theca and granulosa cell function

**DOI:** 10.1101/2023.06.16.545289

**Authors:** Henrietta Philippa Seaward Brain, Christiana Georgiou, Helen D Mason, Suman Rice

## Abstract

Numerous observational and interventional studies have investigated the link between Vitamin D (VD) deficiency and reproductive outcomes, with contradictory results. VD is known to regulate steroidogenic enzymes crucial for human granulosa and cumulus cell function and genes that play a critical role in folliculogenesis have a vitamin D response element (VDRE) on their promoters. This study investigated whether deficient levels of 1,25-(OH)_2_-D3 altered ovarian cell function; and if the ovary could obtain bioactive 1,25-(OH)_2_-D3 via local enzymatic expression of *CYP27B1*, to counteract systemic deficiency. A variety of cells and tissues were used for the *in vitro* experiments as practicable.

We have shown for the first time an increase in VDR expression in <u>theca</u> of larger compared to smaller follicles, which along with the ability of 1,25-(OH)_2_-D3 to decrease Anti-Mullerian hormone (AMH) expression, supports a role for 1,25-(OH)_2_-D3 in theca and granulosa cell function. Conversely, we have shown that very levels of 1,25-(OH)_2_-D3 equivalent to hypovitaminosis, inhibited thecal production of androstenedione and cAMP-driven E2 production. Human thecal and unluteinised GC are incredibly hard to obtain for research purposes, highlighting the uniqueness of our data set. For the first time we have demonstrated that deficient levels of 1,25-(OH)_2_-D3 also down-regulated insulin receptor expression, potentially reducing insulin sensitivity. We have shown that the ovary expresses *CYP27B1* allowing it to make local bioactive 1,25-(OH)_2_-D3 which along with the upregulation in VDR expression in all ovarian cellular compartments, could be protective locally in counteracting systemic VD deficiency. To conclude a severely deficient VD environment (<2nM or <1ng/ml) could contribute to impaired ovarian cell function and hence potentially affect folliculogenesis/ovulation, but levels associated with mild deficiency may have less impact, apart from in the presence of hyperinsulinemia and insulin resistance.

## Introduction

Vitamin D (VD), is a fat soluble prohormone which has both well-established actions such as its classical role in calcium homeostasis and skeletal integrity (Dusso, 2005), and putative ones such asanti-inflammatory and immunosuppressive roles(Aranow, 2011). The discovery of vitamin D receptors (VDR) on male and female gonadal cells prompted investigations into the role of vitamin D in reproduction and fertility (reviewed in Lerchbaum & Obermayer-Pietsch, 2012); (Lorenzen, 2017).

VD is derived minimally from the diet in the form of vitamin D2, with the major source being the photolytic conversion of 7-dehydrocholesterol in the skin to cholecalciferol (vitamin D3) catalysed by UVB radiation (reviewed in Bouillon R *et al*, 2008). Both forms of VD are biologically inert and require activation through sequential hydroxylation in the liver and kidneys. The initial hydroxylation in the liver by 25-hydroxylase enzyme (encoded by *CYP2R1*) results in 25-hydroxyvitamin D3 *aka calcidiol* (25-(OH)-D3), an inert stable metabolite that circulates bound to VD-binding protein (VDBP). The stability of the bound circulating 25-(OH)-D3 means that it is used as an indicator of VD status in an individual (Holick, 2007)). 25-(OH)-D3 is then further metabolised to the biologically active form 1,25-(OH)_2_-D3 (*aka calcitriol*), primarily in the kidneys, a reaction catalysed by the mitochondrial enzyme 1α-hydroxylase (encoded for by *CYP27B1*) (Schuster, 2011); (Luk J, 2012).

1-α-hydroxylase has also been found in extra-renal tissue indicating local conversion of 25-(OH)-D3 to the active 1α,25-(OH)_2_-D3. *CYP27B1* expression has been detected in follicles from macaque and porcine ovaries (Xu J et al, 2018; Grzesiak M et al, 2020) and these studies showed that 1,25-(OH)_2_-D3 regulated follicle growth. Expression of 1,25-(OH)_2_-D3 mRNA and protein was also demonstrated in human granulosa luteal cells (GLCs) and in both benign and malignant ovarian tissue (Fischer D et al, 2009). Thus, it is clear that ovarian expression of 1-α-hydroxylase is capable of producing active 1,25-(OH)_2_-D3 locally within the ovary from circulating precursors.

The ability of active 1α,25-(OH)_2_-D3 to exert its profound actions is mediated exclusively by VDR, a member of the nuclear hormone receptor super-family. In addition to its non-genomic actions outside of the nucleus, VDR acts as a ligand-inducible transcription factor (Haussler MR, 2013), enabling it to regulate 3% of the human genome (Holick, 2007). Normally 1α,25-(OH)_2_-D3 enters the cell by diffusion where it binds to and activates VDR to form a heterodimer complex with retinoid X receptor (RXR): this complex interacts with VD response elements (VDREs) found in the promoter regions of both positively and negatively controlled genes (Cheskis & Freedman, 1994). The localization of VDR to numerous cell types highlights the importance of 1,25-(OH)_2_-D3 as a regulatory hormone in many systems; consequently, VD deficiency is recognised to be associated with numerous pathological conditions which is usually reversed upon supplementation (Holick MF *et al*, 2011;2012).

Current guidelines define VD deficiency as 25-(OH)-D3 levels <20ng/ml (or <50nmol/L); with levels of 21-29ng/ml (52.5-72.5nmol/L) characterized as VD insufficiency and >30ng/ml (>75nmol/L) considered as sufficient (Holick MF *et al*, 2011; 2012) (Pilz S, 2019). Due to avoidance of direct exposure to sunlight and/or nutrient-deficient diets, global deficiency of VD is commonplace in both men and women, but this is exacerbated in northern climates due to lack of sunshine.

The first indication that VD affected female fertility was shown by VD-deficient rats with reduced fertility rates and litter sizes (reviewed in Lorenzen *et al*, 2017). VD has been shown to interact and regulate steroidogenic enzymes that are crucial for human granulosa and cumulus cell function (Merhi Z *et al*, 2014). Interestingly, genes for insulin receptor *(InsR)*, anti-Mϋllerian hormone *(AMH)* and *CYP19A1* (encoding aromatase) that play a critical role in folliculogenesis have VDRE on their promoters (Tiejun S *et al*, 1998); (Maestro B *et al*, 2003); (Krishnan AV *et al*, 2007 & 2010), suggesting a role for VD in female fertility. This is supported by studies showing that *in vitro* supplementation of VD promoted oocyte growth in *in-vitro* developed macaque antral follicles (Xu J et al, 2016). Furthermore, there is a growing body of evidence that VD has a role to play in fertility outcomes in assisted reproductive technology and in women with polycystic ovary syndrome (PCOS), but the results are conflicting (reviewed Lorenzen M. B., 2017).

PCOS is well-recognised as one of the most common endocrine disorders in women, manifesting with increased androgen production, irregular menstrual cycles, and dysregulated ovarian function. (Pellatt L *et al*, 2010). Additionally, there is a clear link between vitamin D, glucose clearance and insulin secretion, such that insulin sensitivity is reduced in vitamin D-deficient women with PCOS, sometimes independently of obesity (Łagowska *et al*, 2018). The compensatory hyper-insulinaemia is known to further amplify the increased thecal androgen secretion in these ovaries (Nestler J.E., 1998) and vitamin D replacement has been shown to reduce serum androgens in some cases (Menichini D., 2020). Interestingly this reduction in hyperandrogenemia can occur in the absence of changes in insulin sensitivity, suggesting a possible direct mechanism of action of VD (reviewed in Menichini D., 2020), though it has to be acknowledged that obesity is a common confounding factor.

Women with PCOS have lower serum 25-(OH)-D3 levels compared to fertile controls (Krul-Poel YHM *et al*, 2018) (Eftekhar M *et al*, 2020) and yet studies investigating the effects of VD deficiency/VD levels on reproductive outcomes produce extremely variable results (reviewed in Lorenzen M. *et al*, 2017). Whilst this could be due to factors such as differing study design, low participant numbers, different ethnicities etc, there is also a lack of understanding of the mechanistic actions of VD in ovarian physiology, and the effect of varying levels of VD ligand interaction with its receptor, which may explain the observed outcomes.

The aim of our study was to investigate the function of ovarian cells in a low vitamin D environment (i.e., ≤20nM), thereby providing mechanistic insight to account for the variable reproductive outcomes observed clinically in women with deficient serum levels of VD. Ideally we would have performed all experiments using human primary ovarian cells and tissues but the scarcity of this source meant we had to use a variety of model systems. In addition, whole ovaries were never given for research, as part were retained for pathology.

Our collaboration with St. Lukes Hospital, Malta gave us access to human primary ovarian cells and tissues, which was a valuable and scarce resource. Follicles were dissected from partial human ovarian specimens to yield granulosa and theca cells for culture, and pieces of cortex and stroma that were snap frozen. The ovarian follicles (preantral and early antral) are in the cortical region. The stroma refers to components of the ovary that are not follicles and include general components such as immune cells, blood vessels, stromal cells and extra-cellular matrix. Over time pre-menopausal oophorectomy was justifiably phased out.

Where feasible we have utilised our stored specimens and material, along with human luteinised granulosa cells (GLCs) from women undergoing IVF for functional experiments. For more complex experiments requiring large number of cells, we have utilised the KGN granulosa cell line. For preliminary immunohistochemistry (IHC) to determine VDR distribution in various ovarian compartments, fixed mouse ovaries were used as no appropriate human samples were available.

## Materials and Methods

All reagents were obtained from Sigma, Poole, UK, unless stated otherwise, and all plasticware was purchased from Fisher Scientific, UK.

### Tissue samples for *in vitro* experiments

A variety of cells and tissues were used for the *in vitro* experiments of this study as detailed below.

#### Human Ovarian Tissues

Informed written consent was obtained from women undergoing trans-abdominal hysterectomy with bilateral oophorectomy for benign gynaecological conditions at St George’s Hospital, London and St Luke’s Hospital, Malta. Ethical approval was granted by the relevant committee in each institution and informed consent obtained. Clinical details were obtained including age, gynaecological history, menstrual frequency, and day of cycle. The ovaries removed from each patient were seen by a pathologist before a portion of each was taken to the laboratory for dissection. Morphology and ovulatory status were assigned as previously published and were based on ovarian size, follicle sizes and numbers, the presence of a dominant follicle or corpus luteum and the amount and density of stroma as determined by dissection, in conjunction with patient history (Mason HD *et al*, 1994; Gilling Smith C *et al*, 1994). The timing of surgery was random. Patient details, number of follicles obtained, and tissue used in each experiment are shown in Table 1. Follicles were isolated from the surrounding stroma, the diameter measured, and granulosa cells (GC) collected as previously described (Mason H. D. *et al*, 1990; 1996). Finally, the theca cell layer was carefully peeled off and digested in an enzyme cocktail for 30mins at 37C with gentle agitation (Mason H. D. *et al*, 1990; 1996). GC and theca cells were cultures as outlined in subsequent sections, other ovarian tissue samples were immersed in RNA-later or flash-frozen and archived at - 80C for further analysis.

#### Luteinised granulosa cells (GLC)

were obtained from follicular aspirates obtained during oocyte retrieval from women undergoing *in-vitro* fertilisation (IVF) treatment (see introduction) at various assisted conception units including those at King’s College Hospital and The Lister Hospital. Local ethics committee approval for each unit was sought and informed consent was obtained from all women.

#### KGN granulosa cell (KGN-GC) line

that are established to correspond to immature granulosa cells from smaller antral follicles, were used to provide mechanistic insight (Nishi *et al*., 2001). The distribution of VDR and AMH protein in various ovarian tissue compartments was assessed using *ovaries from mice (strain 129Sv)* aged between 4 and 5 months. These were euthanised as part of separate ethically approved projects at SGUL and ovaries donated to this study.

### Immunohistochemistry of AMH and VDR in mouse ovarian sections

Ovaries were fixed in paraformaldehyde (4%) at 4C, dehydrated in a gradient ethanol series, cleared in methyl salicylate, embedded in wax and 5μm serial sections cut. Every 3^rd^ section was stained with haematoxylin and eosin (H&E). Adjacent sections were used for immunohistochemistry with anti-VDR (1:100, rat) and anti-AMH (1:50, mouse) monoclonal antibodies (Abcam, Cambridge, UK. Details listed in Table 2) as per manufacturers’ protocol with modifications and appropriate controls.

Briefly, slides were dewaxed and rehydrated, and the antigen epitopes exposed by heating the slides in a 0.01M citrate buffer (pH 6, National Diagnostics) bath until boiling point for 20 minutes. Peroxidases were blocked by 3% hydrogen peroxide (VWR) in methanol (National Diagnostics) for 10 minutes. Sections were then blocked with relevant anti-serum to reduce non-specific binding for 1h at room temperature. Slides were then incubated with the relevant primary antibody overnight at 4C, followed by 1-hour incubation at room temperature with 0.1% monoclonal biotin-labelled secondary antibody (Vector) (Table 2). Visualisation was carried out with Avidin Biotin Complex (ABC) peroxidase solution followed by 3,3’-diaminobenzidine (DAB) (Vector Laboratories). A haematoxylin counterstain was also used for 10 seconds to label nuclei blue. After a brief dehydration through an ethanol series, slides were mounted using Histomount (National Diagnostics). Negative controls omitted primary antibody. Images were captured using the Hamamatsu NanoZoomer 2.0-Rs Slide Scanner for analysis with the NDP.view 2 software (courtesy of the Image Resource Facility, SGUL).

### mRNA and protein expression of VDR in ovarian tissue/cellular compartments

The mRNA expression of VDR in human ovarian cortex, stroma and theca tissue compartments was assessed using archived frozen tissue collected as described above. All tissue was stored and used under HTA regulations. Tissues were defrosted on ice and homogenised in lysis buffer (RLT lysis buffer, Qiagen, Netherlands) using the FastPrep® TissueLyser 24 and the FastPrep® Lysing Matrix A 2mL tubes containing a garnet matrix and a ceramic sphere (MP Biomedicals). After homogenisation RNA was extracted, reverse transcribed and real-time quantitative PCR (qPCR) performed for VDR relative to L19 (the reference gene), as previously described (Rice S *et al*, 2006). In addition, the expression of VDR in the KGN-GC was established using standard PCR as well as qPCR (see Table 3 for details of primers and cycling conditions). For protein extraction the tissue was homogenised in RIPA buffer with a cocktail of phosphatase inhibitors (Sigma-Aldrich, Merck, Gillingham, Dorset), using a sonicator three times for 30s each time. Samples were maintained on ice throughout to prevent overheating and denaturation of proteins. The lysed samples were micro-centrifuged at 4C for 10min at maximum speed. Pelleted debris was discarded, and the protein lysate stored at −80C prior to Bradford assays (for measurement of protein concentration) and western blotting.

### GC, theca and GLC culture experiments

The granulosa or theca cells were pooled from several follicles based on the experimental protocol and cultured in McCoys 5A medium supplemented with penicillin/streptomycin, L-glutamine and amphotericin B (all purchased from Invitrogen, UK). Granulosa cells were plated in 96-well plates at 5×10^4^ and theca cells at 5×10^5^ cells per well as previously described (Mason *et al*, 1990; Willis *et al*, 1996). To increase the adhesion of theca cells to the plate, plates were first coated in ECM gel (McCaig, 2002).

Cells were incubated in culture medium (plus experimental treatments) for 48 hours with a range of 1,25-(OH)_2_-D3 concentrations (0.2, 2 and 20nM serially diluted down from a stock concentration of 2µM). Testosterone (5× 10^−7^ M) was used as a substrate for conversion to oestradiol in the granulosa cells and steroid levels measured in the medium by radioimmunoassay (Hillier SG *et al*, 1980); (Gilling-Smith C *et al*, 1994); (Willis DS *et al*, 1996). To confirm that cells were healthy, 10ng/ml LH was used as a positive control in luteinised GCs and theca, and 5ng/ml FSH in cells dissected from small follicles (LH and FSH supplied by Endocrine Services, Biddeford-upon-Avon, UK). Isolation of GLC was performed as previously described (Wright R *et al*, 2002). Cells were plated in 96-wells at 5×10^4^ cells per well in 200μl of serum (5%) supplemented medium for 48 hours, after which the medium was removed and replaced with experimental treatment as detailed above. GLCs were also exposed to the vitamin D analogue ED1089 (kindly donated by Dr. Kay Colston, SGUL), to examine comparative efficacy and demonstrate specificity.

### KGN-GC culture experiments

To model the effect that VD deficiency may have on ovarian function, KGN-GC were grown and passaged in 10% DMEM-F12 supplemented with L-glutamine and penicillin/streptomycin (Invitrogen), at 37◦C in 95% air/CO_2_. Cells were plated in 12-well plates (3×10^3^ cells/well) and cultured in 1% DMEM-F12 (charcoal-stripped) overnight. Cells were serum-starved for 2h and then treated with forskolin (25μM), to reproduce the effect of LH on cAMP stimulation, ± 1,25-(OH)_2_-D3 at 0.02, 0.2, 2 and 20nM and cultured for a further 48h. Testosterone (5×10^−7^M) was added to all cultures as the aromatase substrate for conversion to oestrogen. Similarly, the effect of VD on AMH mRNA expression was investigated in cells treated with forskolin and VD as described above, using only the upper and lower doses of 1,25-(OH)_2_-D3 from the range above at 0.02 and 20nM. At the end of the relevant culture time, RNA was extracted, reverse transcribed and real-time quantitative PCR (qPCR) performed for *CYP19A1* (encoding aromatase) as described previously (Rice S *et al*, 2013). We have previously established that *CYP19A1/aromatase* mRNA, protein expression and enzyme activity are all tightly correlated in this model and hence any of these techniques can be used interchangeably to assess changes (Rice *et al*, 2006; Rice *et al*, 2013).

To investigate the effect of VD on aromatase promoter II (PII) activity, cells were plated at a density of 2×10^4^ cells/well in 96-well plates. After overnight incubation, they were transfected with a *CYP19A1 PII-516* reporter construct expressing firefly Luciferase, along with a control plasmid expressing Renilla luciferase from a constitutive promoter, and a transcription enhancing element, PVAi, as previously described (Rice S *et al*, 2013). After 2h serum-starvation, cells were treated as described above for 24 hours with quadruplicate replicates/treatment, and luciferase reporter assays were carried out using the Dual-Glo Luciferase Assay System (Promega).

To mimic a situation of chronic insulin exposure, cells were treated with insulin for 48h at post-prandial (10ng/ml) and hyperinsulinemic (100ng/ml) levels in the presence and absence of the lowest [0.02nM] and highest [20nM] 1,25-(OH)_2_-D3 levels tested in the experiments. The effect of 1,25-(OH)_2_-D3 on aromatase and insulin receptor (InsR) mRNA expression was measured by qPCR.

### Radioimmunoassay (RIA) for P, E2 and Androstenedione

Conditioned media was collected from the cells at 48 hours, frozen at −20C and E2, P and androstenedione measured using a modified ‘in-house’ RIA, with tritiated steroids and charcoal separation (Hillier SG *et al*, 1980); (Gilling-Smith C *et al*, 1994); (Willis DS *et al*, 1996). The components of the assay were: tritiated steroid label (Amersham Pharmaceuticals, Bucks, UK), steroid antiserum (sheep anti-human from Guildhay, Guildford, UK) and standards and quality controls (QCs) (approximately 1, 6 and 35% of the top standard), prepared from powdered steroid (Sigma Co Ltd, Poole, UK). Conditioned medium was diluted to be within the midrange values for each assay and therefore on the linear part of the standard curve. Each sample was tested in duplicate.

### Immunoprecipitation (IP)

To investigate the effect of 1,25-(OH)_2_-D3 on VDR:RXR dimer formation, KGN cells were plated at a density of 5×10^5^ cells/well in a 6-well plate and treated as described above with and without forskolin, testosterone and 1,25-(OH)_2_-D3 at 0.02 and 20nM for 48h. At the end of culture time, media was removed, cells washed with ice-cold PBS and lysed with ice-cold RIPA buffer as previously described (Dilaver N *et al*, 2019). The protein concentration of the lysate was measured by Bradford assay and prior to immunoprecipitation, equal amounts of protein lysate from each treatment group were pre-cleared on A/G-coupled Sepharose beaded (Pierce) support (binding capacity 27mg mouse IgG or ≥ human IgG/ml settled resin) for one hour at 4C to reduce non-specific ligand binding. Immunoprecipitation of the cleared lysates was performed overnight at 4C with 5μg of anti-VDR antibody or anti-RXRα monoclonal antibodies (Abcam – Table 2). The beaded complexes were washed several times with NP-40 lysis buffer, after which protein complexes were dissociated by boiling for 10 mins with 1X SDS-DTT reducing buffer (protocol adapted from Luderer HF *et al*, 2011). The collected supernatant was stored at −80C for subsequent Western blotting analysis. Negative controls included retention of pre-cleared beads which were eluted and run on western blots, alongside the total lysates.

### Western Blotting of VDR and RXR

Protein concentrations were measured using the Bradford assay (BioRad, Hertfordshire) and equal amounts of protein obtained from lysing ovarian tissues or from immunoprecipitation experiments, were resolved with Western blotting as previously described (Dilaver N *et al*, 2019). PVDF(fl) (Immobilon, Millipore Merk, Massachusetts) membranes with transferred proteins were incubated with rat anti-human VDR or rabbit anti-human RXR (both used at 1:1000) and/or mouse anti-human β-actin (1:2000). Fluorescently conjugated relevant secondary antibodies (1:5000) (see Table 2 for details of antibodies) were used for visualisation using the Odyssey Imaging System (Li-Cor Biosciences) (Dilaver N *et al*, 2019).

### Experimental Quantification and Statistical Analysis

All data are represented as the mean ± SEM of triplicate or more observations (detail in legends) from a minimum of 3 or more independent experiments, unless otherwise stated. qPCR data were analysed using the ΔΔCt method as described in detail previously (Rice S *et al*, 2006), with normalisation to L19 and subsequent normalisation to the Ct value of the control (untreated). To use the ΔΔCt method, the amplification efficiency for each GOI and reference gene must be in the recommended range of 90-100%. This was rigorously applied to our study by the inclusion of a standard curve for every qPCR assay conducted. Data from Western blots represent the mean densitometry measurements taken from all individual experiments using Image Studio software (Licor™) and normalised to ß-actin (loading control) and where relevant, to the untreated (control) samples. For the IP experiments, the densitometry values were also adjusted for the quantity of protein in the sample. Results for steroid RIA were calculated using Assay Zap V2.69 software (Biosoft, Cambridge, UK). Intra-and inter-assay coefficients of variation were below 5% and 6% respectively.

Statistical significance was determined by ANOVA followed by post hoc multiple comparison tests; unpaired Student’s or paired t test when 2 groups were compared (depending on the design of the experiment) or a one-sample t test when comparing with normalised control values, using GraphPad Prism™. Significance was set at P ≤ 0.05.

## Results

We analysed expression (via qPCR, IHC and WB) of key elements (VDR and *CYP27B1)* involved in the synthesis and signalling pathway of VD in various ovarian cellular compartments. We then proceeded to investigate the effect of a range of doses of VD on steroid output (RIA) from primary theca and unluteinised granulosa cells dissected from antral follicles of varying sizes cultured *in vitro*. The effect of VD on oestradiol (E2) production (RIA and qPCR) was also analysed in luteinised GLCs and the KGN cell-line. Since VD is known to interact with other factors involved in granulosa cell proliferation, we chose two of particular interest, namely AMH and insulin to investigate via qPCR. Finally, we examined the relationship via IP between varying doses of VD and the degree of binding to VDR and RXR receptors, which then complex to VDREs to modulate its actions.

### Expression of VDR and *CYP27B1* in ovarian tissue and cellular compartments

IHC was used to detect the extent of VDR protein expression in sequential mouse ovary sections, and the anti-VDR antibody revealed strong and extensive staining throughout all mouse ovarian tissue compartments including the cellular components of the follicles (figure 1a (negative control) and 1b). Closer examination revealed more intense staining in thecal cells of LAF compared to SAF (inset figures 1b). The general uniformity and ubiquity of VDR expression was in marked contrast to staining with anti-AMH antibody in the sequential section, which showed a distinctive pattern of expression i.e., greater staining in GC of SAF compared to LAF, with no staining in theca or ovarian compartments (figure 1c).

This finding in the mouse ovary was extended by measuring VDR mRNA levels in human theca cells dissected from human follicles of varying sizes to ascertain whether expression levels changed with follicle size. VDR mRNA levels were significantly higher in theca from larger antral follicles (LAF) and those approaching preovulatory sizes (7-12mm, n=11), compared to those from smaller antral follicles (SAF) (5-6mm, n=4) (unpaired t-test, *p=0.013; figure 2a) which supported the observations in the mouse ovary (fig 1b).

**Fig 1a:**
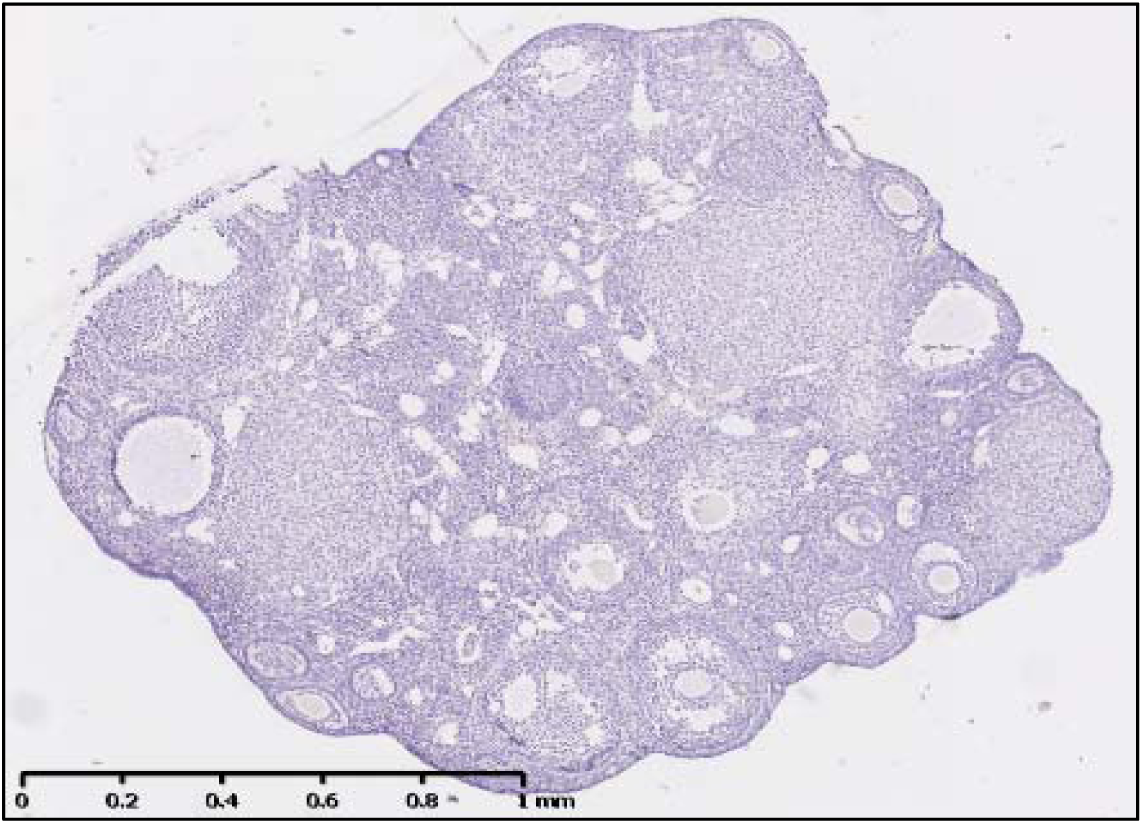
Negative control (no VDR primary antibody) of the sectioned mouse ovary with a variety of small antral follicles (SAF) and large antral follicles LAF).

**Fig 1b:**
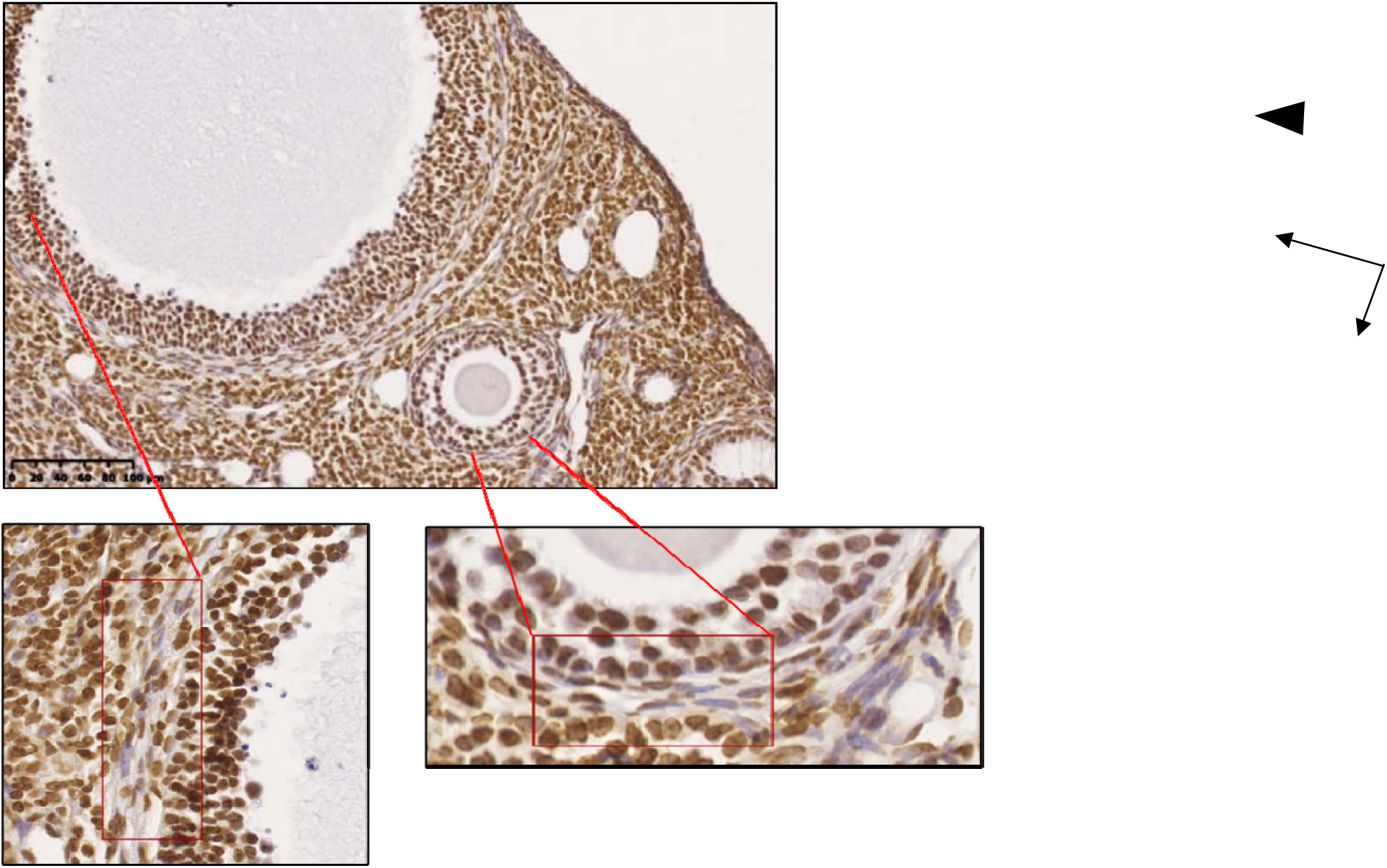
IHC of VDR expression in the corresponding adjacent section of the same mouse ovary. There is extensive expression of VDR protein throughout cortex, stroma and cellular compartments i.e. both GC (arrowheads) and theca (arrows) of both SAF and LAF. Though the intensity of staining in theca appears to be greater in theca of LAF compared to SAF *(inset pictures enlarged)*.

**Fig 1c:**
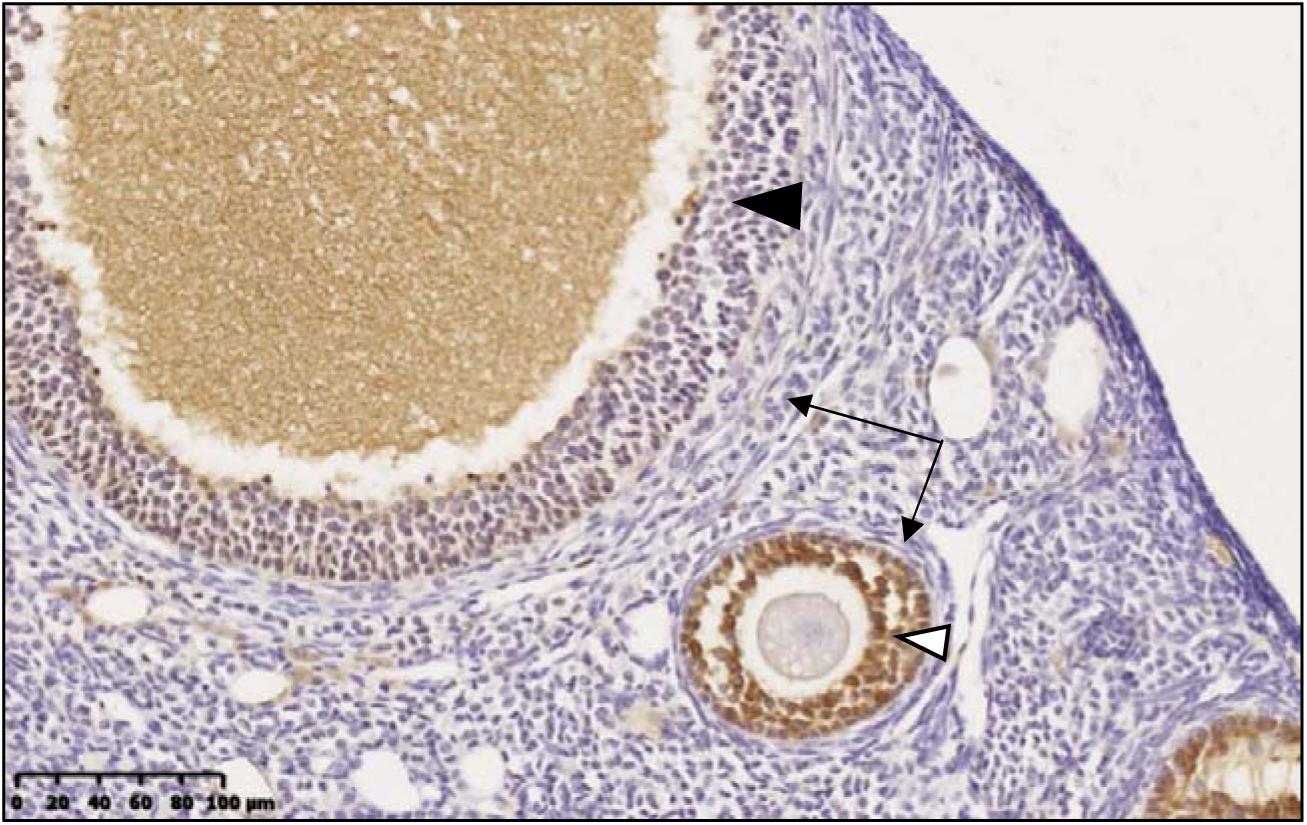
IHC of AMH expression in the corresponding adjacent section of mouse ovary. AMH protein is expressed strongly in GC of SAF (white arrowhead) with minimal expression in GC of LAF (black arrowhead). There is <u>no expression</u> in theca cells (black arrows) or in cortex/stroma.

**Fig 2a:**
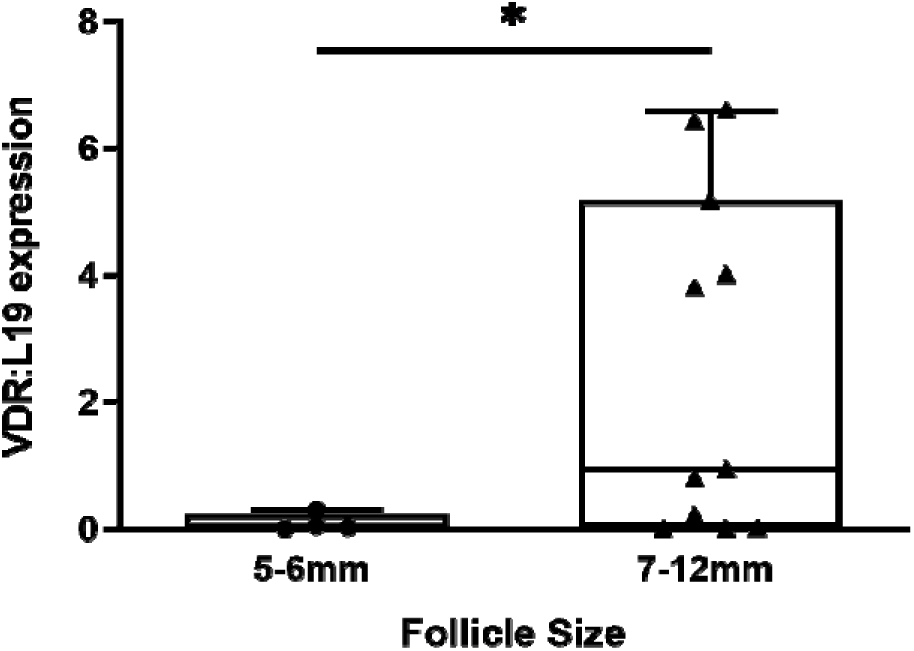
mRNA expression of VDR quantified by qPCR, in <u>human theca</u> taken from human ovarian follicles (both normal and polycystic ovaries), showed significantly lower expression in SAF (5-6mm ●) compared to LAF (7-12mm ▲). *(Unpaired t-test, two-tailed*p=0.0128; 5-6mm (n=4); 7-12mm (n=11))*.

In addition, VDR mRNA (fig 2b) and protein (fig 2c) expression were also detected in human cortex and stroma… There was no statistically significant difference between the two compartments. *CYP27B1* mRNA was expressed in human ovarian stroma as assessed by qPCR, though there was considerable variation in expression levels between different biological samples (fig 2d).

**Fig 2b:**
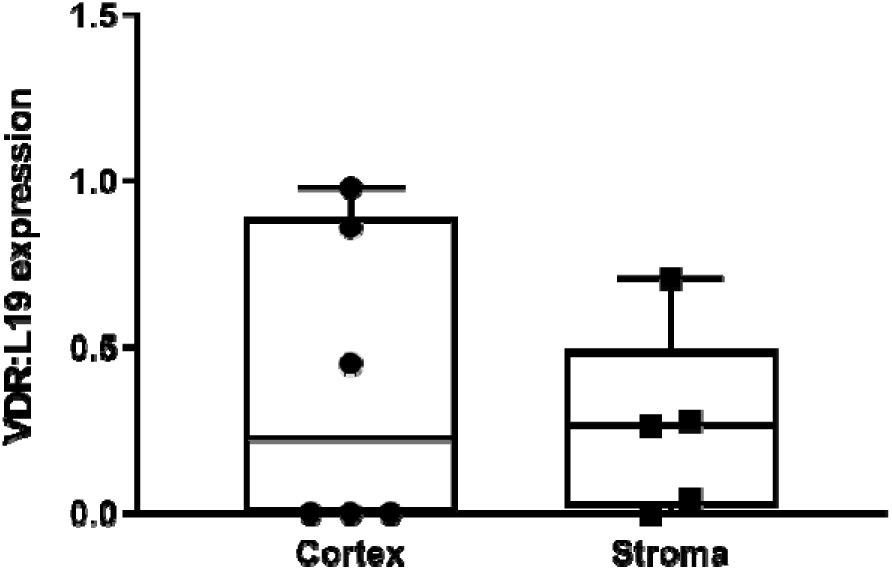
VDR mRNA expression quantified by qPCR in human cortex (grey bar ●, *n=6*) and stroma (white bar ■, *n=5*). There was no significant difference in expression.

**Fig 2c:**
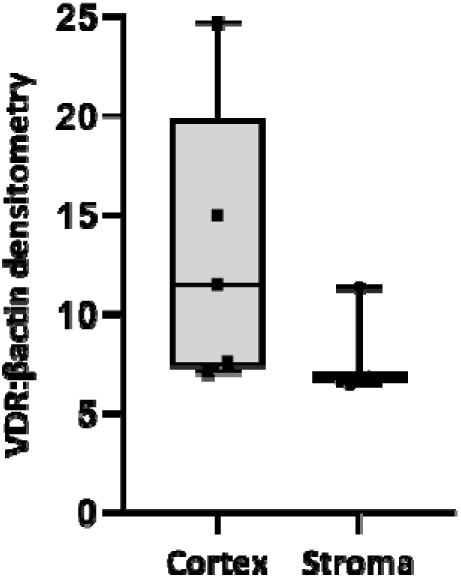
VDR protein expression levels quantified by western blotting, in human ovarian cortex (*n=5; grey bar* ▪) and stroma (*n=4; white bar* ●), also showed no significant difference in expression. *Densitometry values on y-axis multiplied by 1000 for scaling purposes*.

**Fig 2d:**
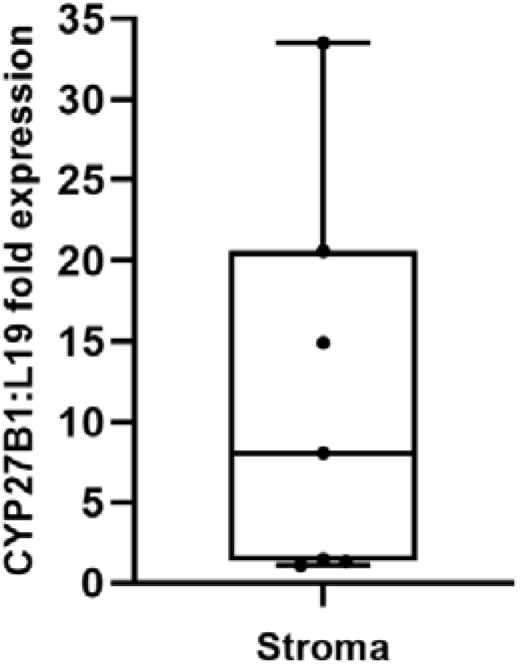
CYP27B1 mRNA expression quantified by qPCR, in human ovarian stroma samples – both normal and PCO *(n=8)*.

### Effect of 1,25 (OH)2-D3 on steroid production in ovarian cells

#### VD and theca steroidogenesis

Overall, in theca from LAF (15-22mm), exposure to all doses of 1,25-(OH)_2_-D3 consistently suppressed Androstenedione production (mean 22% suppression) but had no effect on androstenedione production from SAF (n=4-8, Two-way ANOVA, p=0.048 for treatment and p=0.002 for follicle size) (fig 3a). Treatment with 1,25-(OH)_2_-D3 had no effect on either P or 17-OH-P production (supplemental figures S1 and S2).

**Fig 3a:**
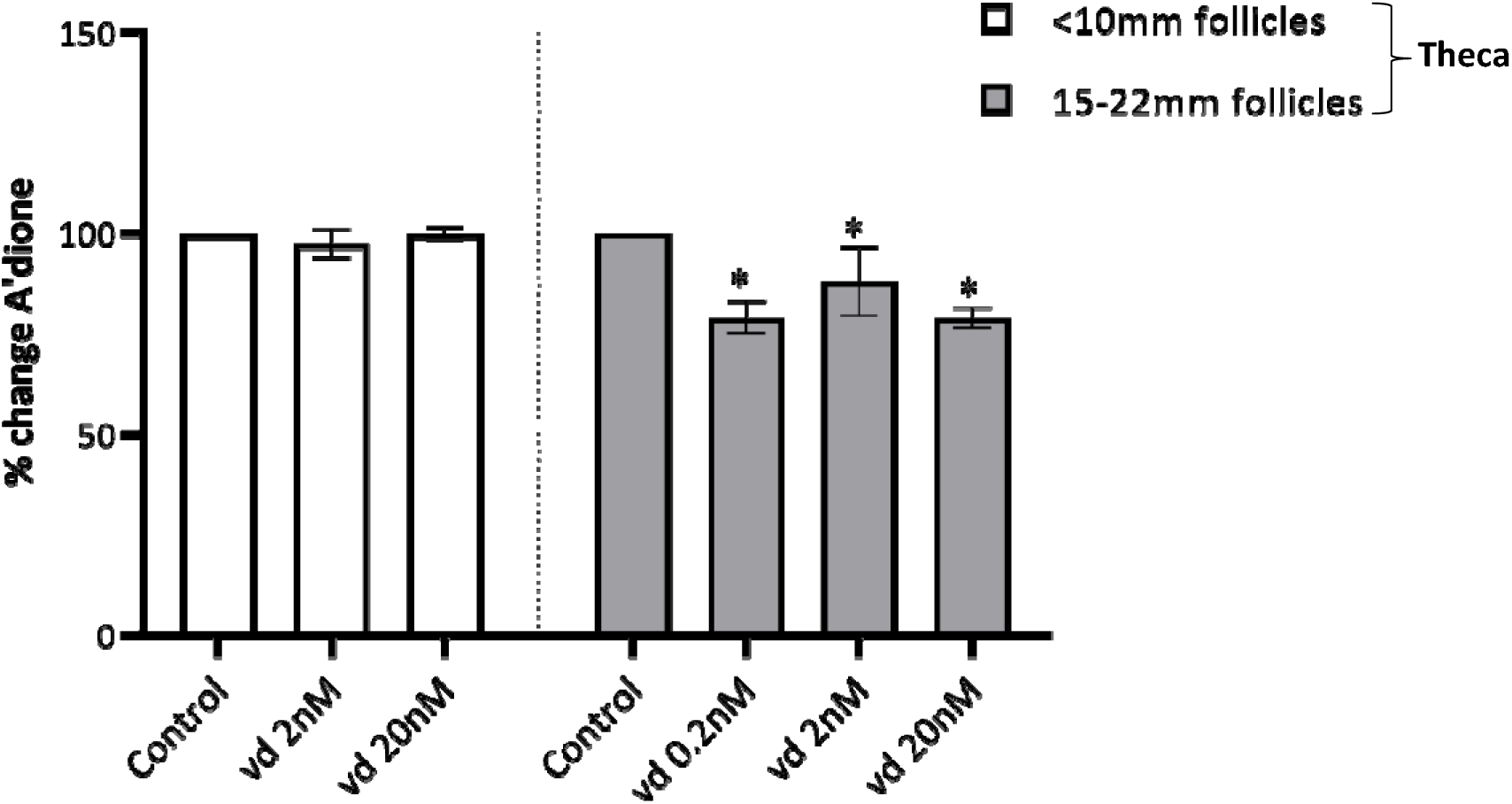
Effect of 1,25(OH)_2_D_3_ on androstenedione production from <u>theca</u> taken from small (<10mm, white bars, *n=4 subjects, total follicles=15*) and large follicles (15-22mm, grey bars, *n=5 subjects, total follicles=5*) from normal ovaries. 1,25(OH)_2_D_3_ suppressed androstenedione production significantly but only in theca from large follicles. Results expressed as a percentage change from control, where the control was taken as 100%. *(Two-way ANOVA p=0.05, source of variation follicle size **p=0.002, treatment*p=0.048; Tukeys post-hoc multiple comparison *p<0.05)*.

#### VD and unluteinised GC steroidogenesis

There was no effect of 1,25-(OH)_2_-D3 at any dose on E2 production in GCs taken from follicles <10mm diameter (n=4; figure 3b); however, in follicles >10mm there appeared to be a trend to increasing E2 with 1,25-(OH)_2_-D3 though this was not significant (figure 3b). In addition, 1,25-(OH)_2_-D3 did not alter the normal responsiveness to FSH in SAF (n=4, ANOVA p<0.0001) (figure 3c).

**Fig 3b:**
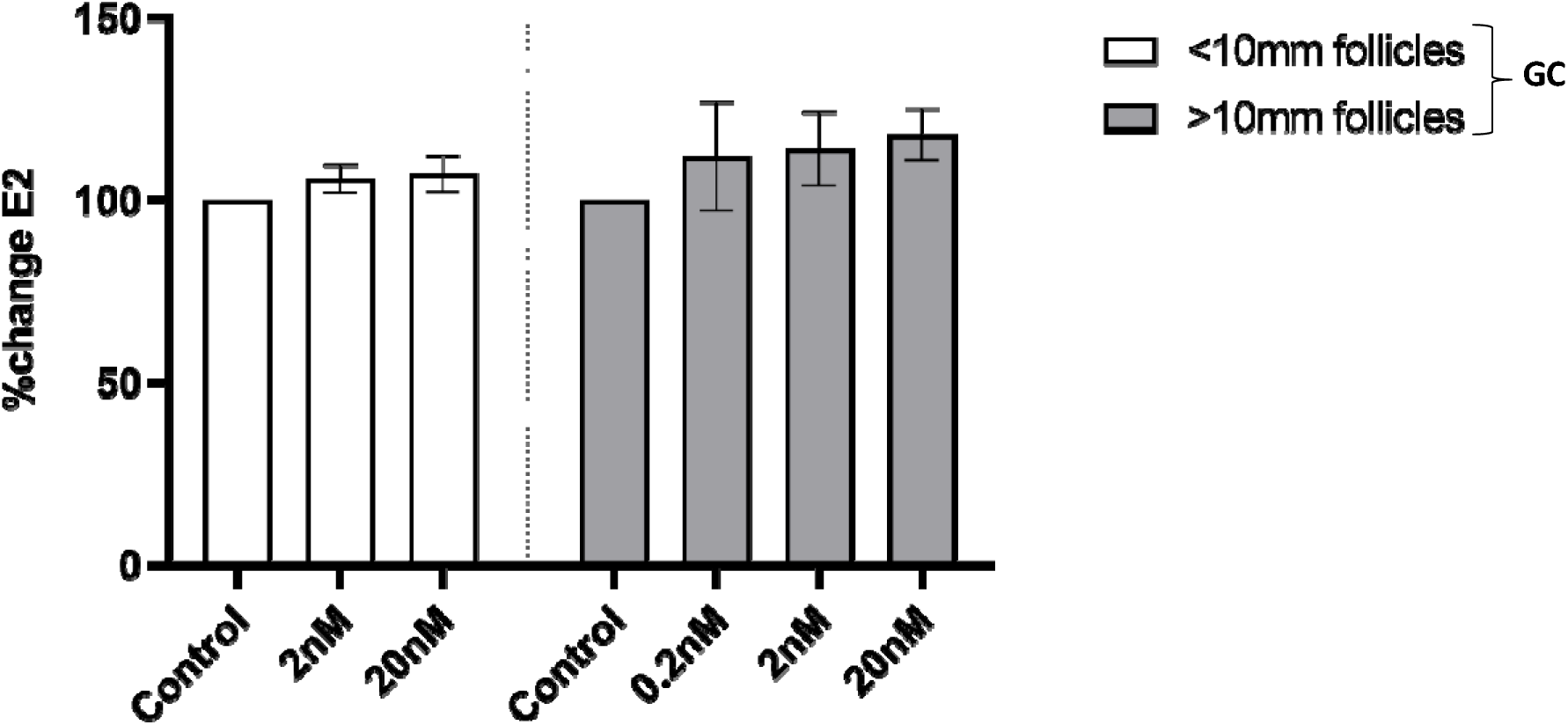
1,25(OH)_2_D_3_ had no effect on E_2_ production from <u>granulosa cells</u> taken from small antral follicles (<10mm, white bars; *n=3 subjects, total follicles=11*) and large antral follciles (>10mm, grey bars; *n=6 subjects, total follicles=8*) from normal ovaries. Results showed no effect of 1,25(OH)_2_D_3_ (vd 0.2-20nM) on E_2_ production. The results are expressed as mean percentage change from control where the control is taken as 100%.

**Fig 3c:**
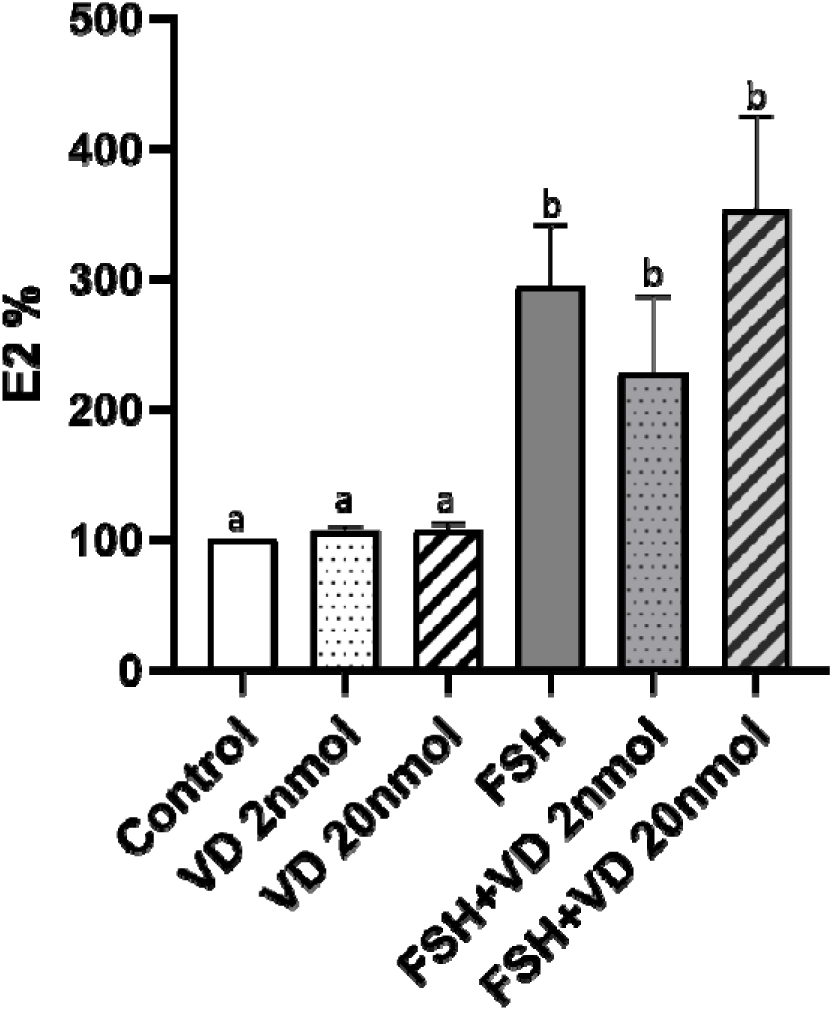
1,25(OH)_2_D_3_ at 2 (spotted bar) & 20nM (hashed bar) had no effect on E_2_ production from <u>granulosa cells</u> in the absence (white bars)/presence(dark grey bars) of FSH (5ng/ml), which as expected significantly simulated E2 production. The results are expressed as mean percentage change from control where the control is taken as 100%. *(n=4 subjects; total follicles=49. ANOVA ****p<0.0001; **p<0.005 Sidak’s multiple comparison test)*.

#### VD and E2 production in GLC

The effect of VD on E2 production from luteinised granulosa cell culture was then investigated. Luteinisation of GC occurs once follicles have been exposed to the mid-cycle ovulatory LH surge. As expected, exposure of the cells to LH produced a substantial stimulation in the production of E2, surprisingly this was considerably attenuated by 1,25-(OH)_2_-D3 at both 2nM and 20nM doses (figure 3d) (ANOVA p<0.0001; post-hoc t-test **p<0.005). This contrasts with the results from unluteinised GC from small follicles in which 1,25-(OH)_2_-D3 had no effect on FSH-stimulated E2 production (figure 3c). Interestingly culturing GLC with the VD analogue EB1089 also appeared to suppress E2 production in a dose-related manner (figure 3e), though the data did not reach statistical significance (ANOVA n/s). There was, however, no effect on progesterone production in the cells by any dose of VD (supplemental figure S3).

**Fig 3d:**
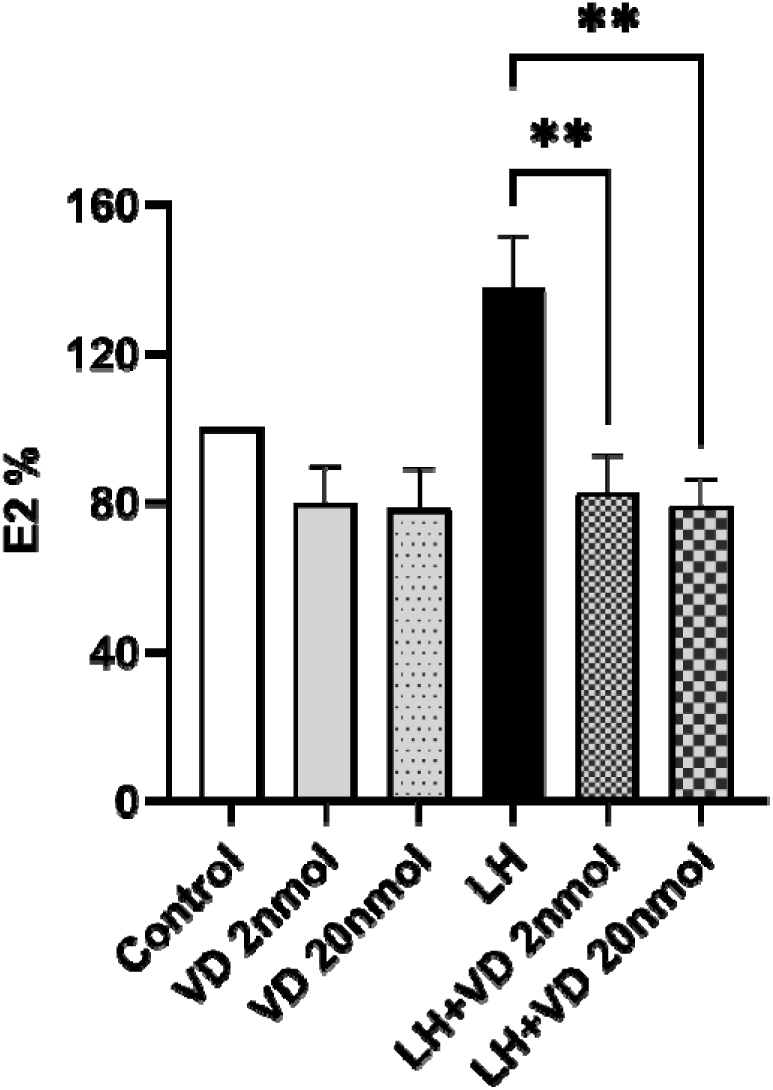
Effect of 1,25(OH)_2_D_3_ (at 2 & 20nM) on E_2_ production from <u>granulosa-lutein cells</u> in the presence of LH (10ng/ml). LH significantly stimulated E2 production, which was significantly attenuated in the presence of VD *(n=4; ANOVA **p<0.005; Post-hock Sidak’s multiple comparison test. Alphabetical annotations are used to denote differences in statistical significance)*.

**Fig 3e:**
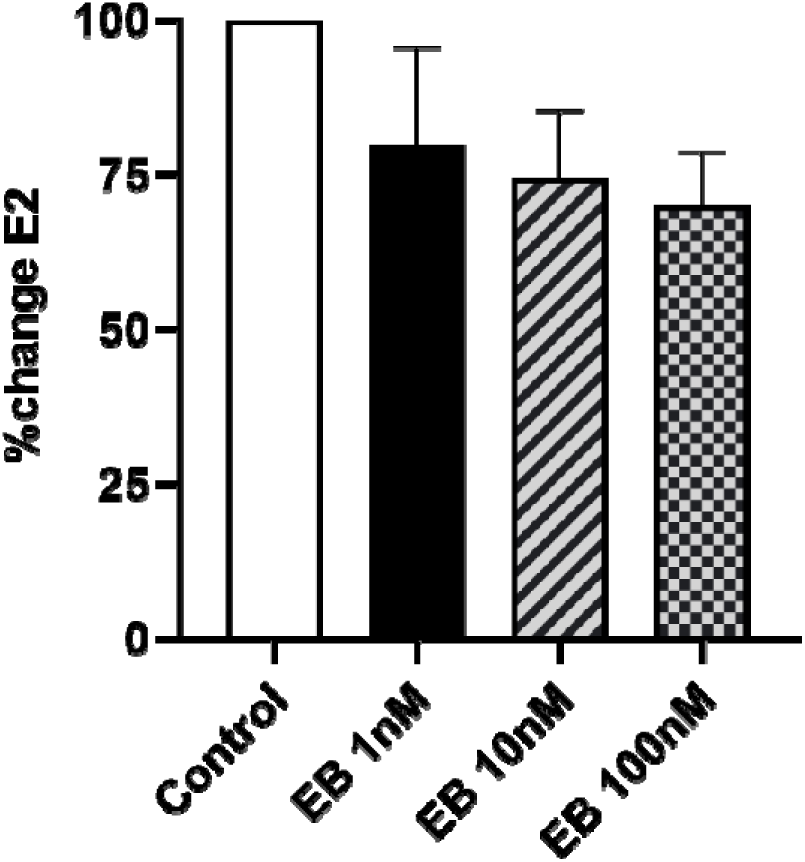
A similar dose-dependent reduction in E2 production was seen in <u>granulosa-lutein cells</u> cultured in the presence of the VD analogue EB1089, though this did not reach statistical significance (*n=3*).

##### Effect of VD on aromatase expression in KGN-GC

To determine the mechanism by which 1,25-(OH)_2_-D3 was altering E2 production in primary cells, KGN cells were cultured with forskolin to increase cAMP and hence aromatase activity (equivalent to stimulation by either gonadotrophin, which both act via cAMP). Very low 1,25-(OH)_2_-D3 levels [0.02 & 0.2nM] significantly down-regulated Fsk-stimulated aromatase mRNA expression (n=6, One-way ANOVA**p=0.0018; post-hoc t-tests *p<0.05, **p<0.005), but when doses approached sufficiency levels [2 and 20nM] this attenuation was reversed. An identical pattern was seen in PII transfected KGN cells demonstrating that 1,25-(OH)_2_-D3 affected aromatase activity as well as expression (figures 4a & 4b) (n=3; ANOVA****p<0.0001; post-hoc t-tests *p=0.02, **p=0.0095).

**Fig 4a:**
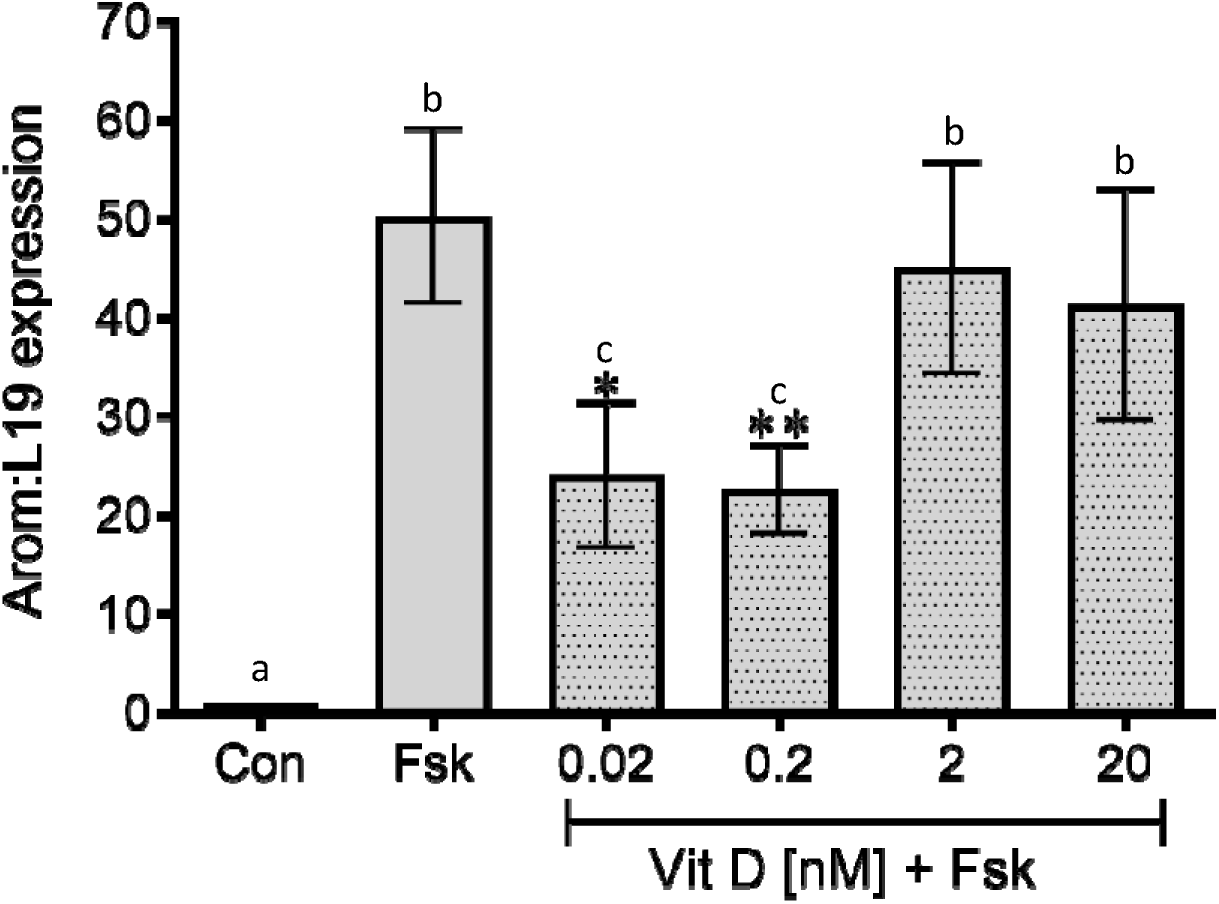
Aromatase mRNA expression levels were measured in KGNs cultured with forskolin (Fsk) to stimulate aromatase and different doses of VD (0.02-20nM). Testosterone (5×10^−7^M) was used as an aromatase substrate. VD at the lowest two doses significantly down-regulated Fsk-stimulated aromatase expression, but as the doses increased this attenuation was lost. *(One-way ANOVA ***p=0.0002; Tukey’s multiple comparisons test con vs fsk **p<0.005; con vs 2 **p<0.005; con vs 20 *p<0.05; fsk vs VD0.02 *p<0.05; fsk vs VD0.2*p<0.05 (n=6). Alphabetical annotations are used to denote differences in statistical significance)*.

**Fig 4b:**
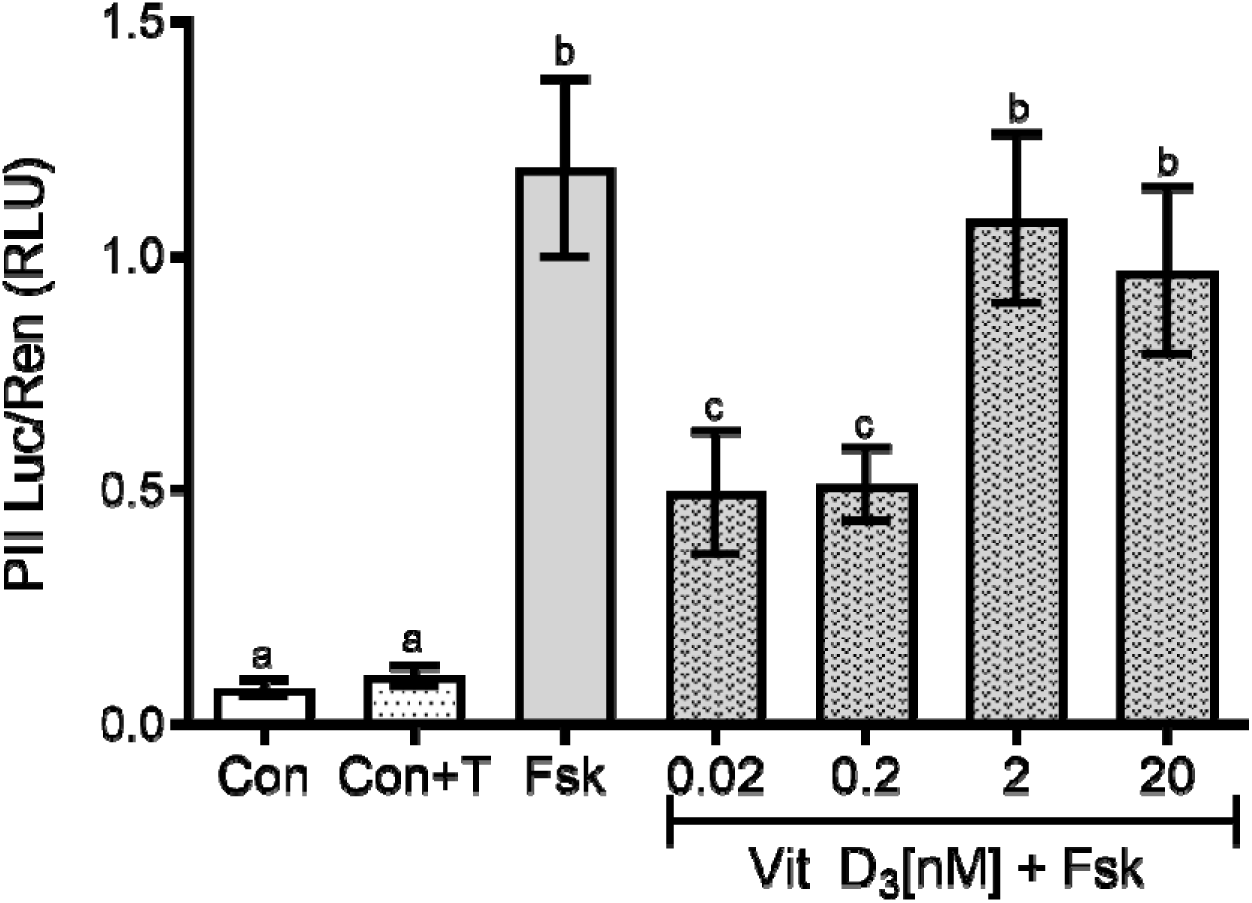
KGN cells were transfected with PII-specific-Luc reporter construct and treated as for the qPCR experiments. The luciferase assay showed that the effect of VD doses on aromatase expression was direct action on aromatase transcription, with very low doses down-regulating Fsk-stimulated PII activity. As VD concentrations increased this attenuation was lost. *(One-way ANOVA****p<0.0001; Tukey’s multiple comparison-C/C+T vs Fsk***; Con vs 2***; Con vs 20***; Fsk vs 0.02**; Fsk vs 0.2*; 0.02 vs 2* (n=3). Alphabetical annotations are used to denote differences in statistical significance)*.

### Effect of VD on other factors implicated in regulation of follicle growth

#### Insulin

There is compelling evidence that systemic VD levels are correlated with insulin sensitivity (Łagowska *et al*, 2018); however, there is little supportive evidence at a cellular level. Chronic exposure of granulosa cells to high (100ng/ml) insulin significantly downregulated expression of total InsR mRNA (p=0.001), and this reduction was surprisingly potentiated by both doses of 1,25-(OH)_2_-D3: 20nM (<50% of basal) and 0.02nM (<25% of basal) (n=5-8; One-way ANOVA *p=0.048, one-sample t-tests to the control ***p<0.0005, **p<0.005, *p<0.05) (figure 5a). This profound attenuation of insulin receptor expression by 1,25-(OH)_2_-D3 in the presence of chronic insulin exposure did not have a concomitant reduction on aromatase expression in the same cells. In fact, the presence of insulin reversed the attenuation of basal aromatase levels brought about by low dose VD (figure 5b).

**Fig 5a:**
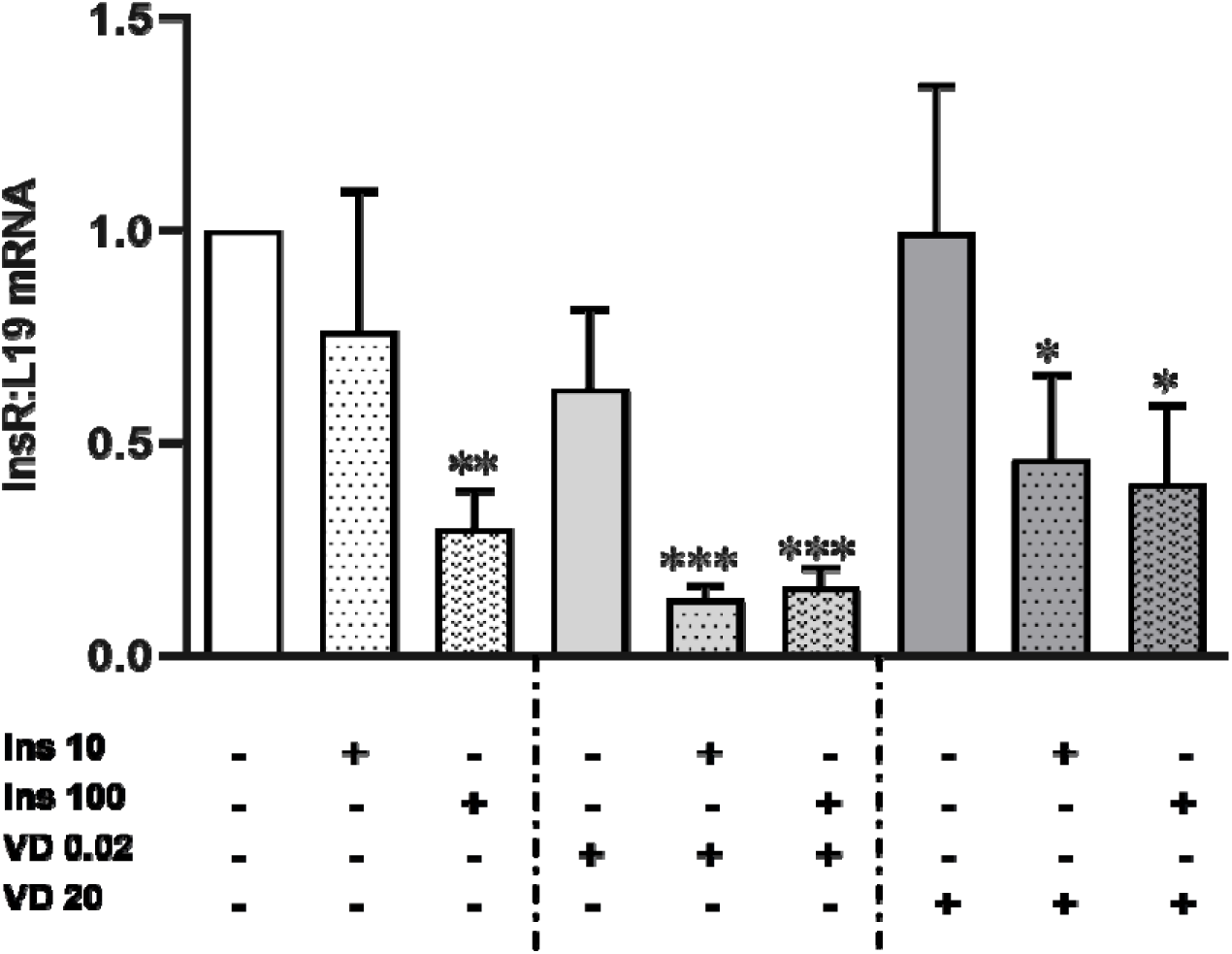
Expression of <u>Insulin receptor</u> (InsR) mRNA expression ± insulin and VD at various doses in KGNs. Chronic exposure to high insulin (100ng/ml) significantly reduced InsR expression (spotted white bars), which was further attenuated in the presence of very low VD (light grey spotted bars). This combined suppression of InsR was seen even with VD at sufficient levels compared to basal (dark grey spotted bars). *One column t-test p=0.0014 Ins100; p=0.001 VD20+Ins10; p=0.0003 VD0.02+Ins100; p=0.0522 VD20+Ins10; p=0.0318 VD20+Ins100. One-way ANOVA*p=0.048; (n=5-8). Values expressed as mean±SEM*.

**Fig 5b:**
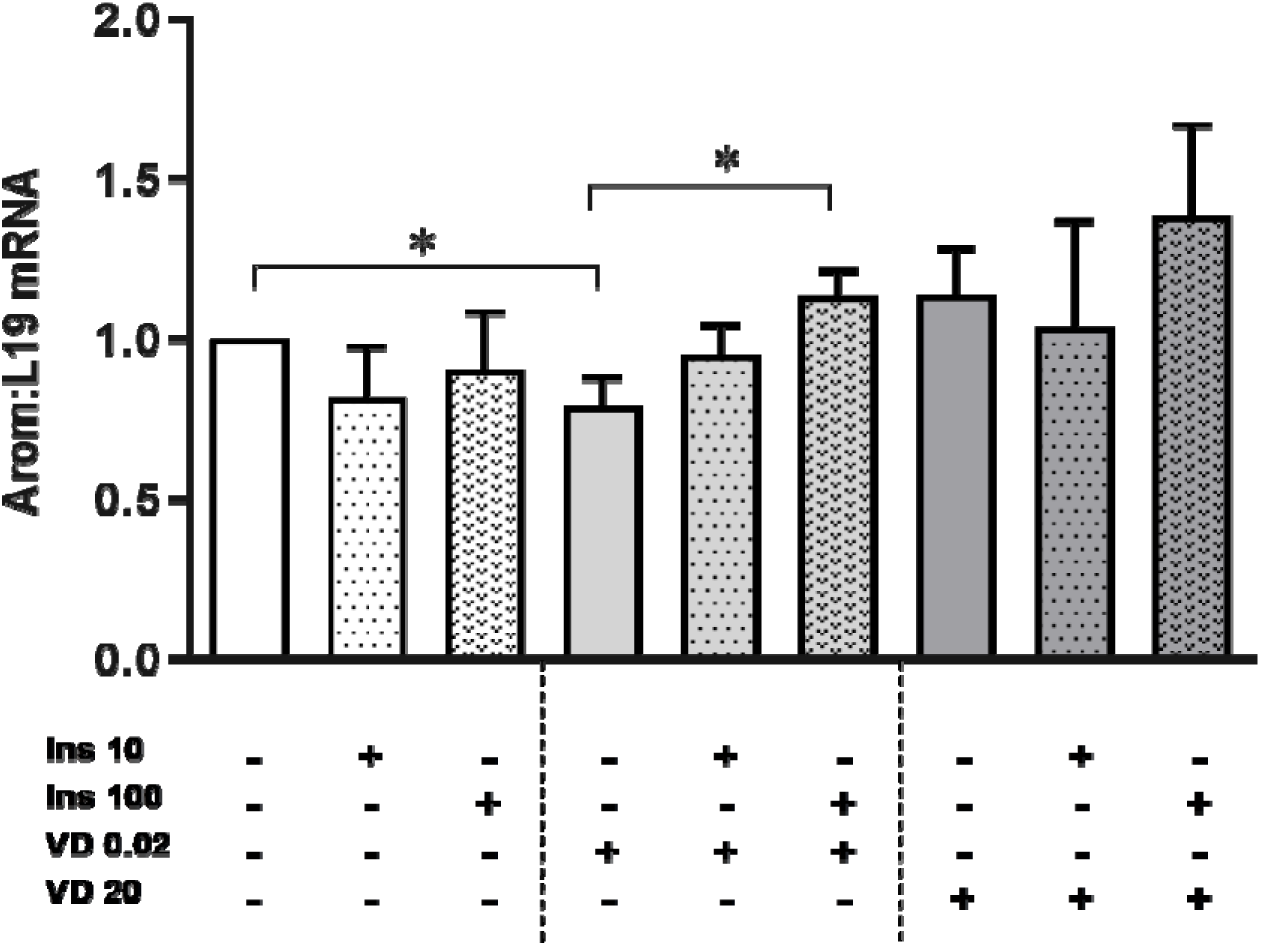
Expression of <u>aromatase</u> mRNA expression ± insulin and VD at various doses in KGNs. Insulin had no effect on aromatase expression but reversed the attenuation of basal aromatase expression brought about by low dose VD exposure (light grey spotted bars). *One way ANOVA *p=0.0307; Unpaired t-test: VD0.02 vs VD0.02+Ins100 *p=0.0118 (2 tail); VD0.02 vs VD20 +p=0.0324 (1-tail) (n=5-8) Values expressed as mean ± SEM)*.

#### AMH

Forskolin down-regulated AMH expression compared to basal (<50%), which was further potentiated in the presence of 1,25-(OH)_2_-D3, with a more potent attenuation by 20nM 1,25-(OH)_2_-D3 compared to 0.02nM (figure 5c) (n=3, ANOVA *p=0.02; multiple comparison t-test *p<0.05).

**Fig 5c:**
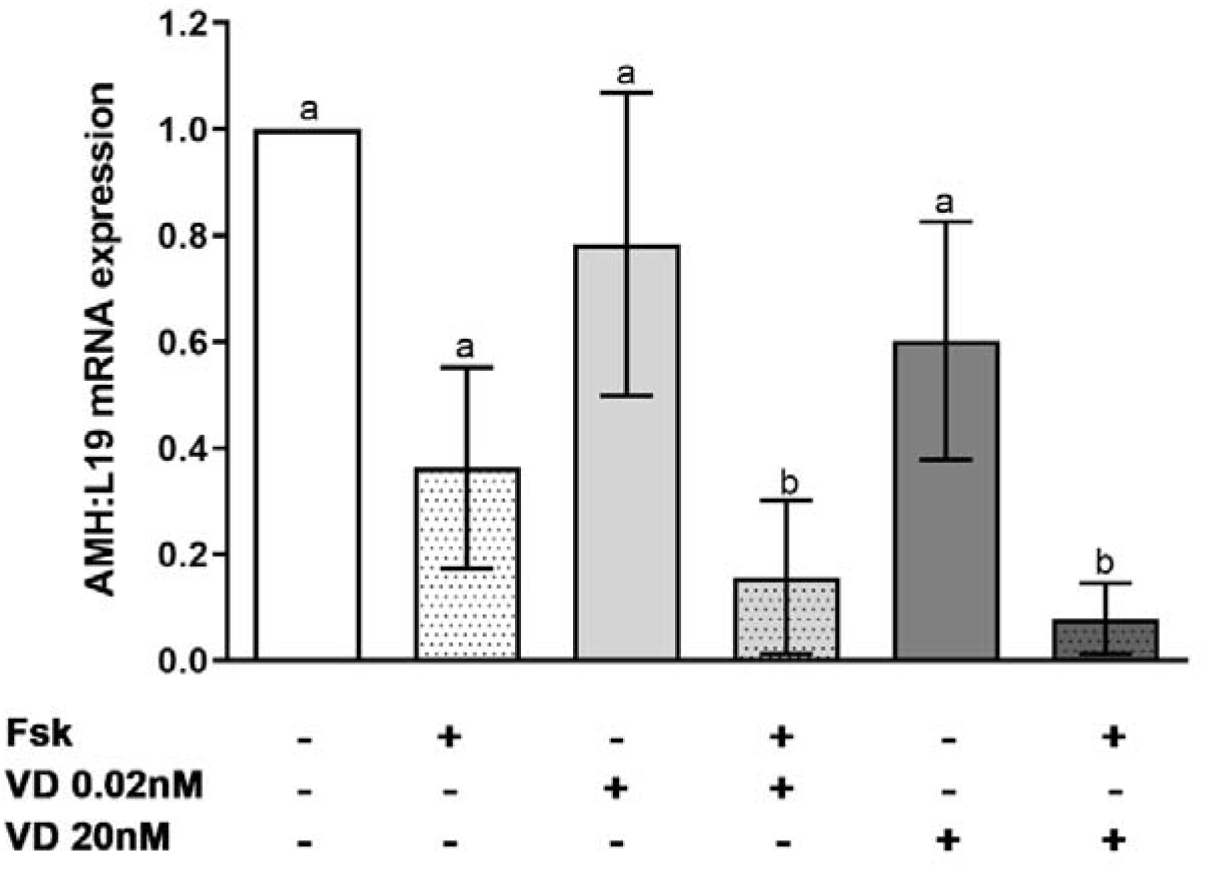
mRNA expression of AMH in the presence of VD (at 0.02 and 20nM) and forskolin (Fsk) in KGNs. Fsk reduced AMH expression below basal (white spotted bars) and this was further attenuated in the presence of 1,25(OH)_2_D_3_ with the strongest reduction seen in the presence of 20nM VD (dark grey spotted bars) *(ANOVA*p=0.02; Tukey’s multiple comparison *p<0.05)*.

### Investigating the differential effects of varying doses of VD ligand

Data presented so far indicate that those doses of 1,25-(OH)_2_-D3 equivalent to extremely deficient levels have differential effects compared to those approaching sufficiency. To determine a possible mechanism, the relationship between levels of VD ligand and the proportion of either VDR or RXR was investigated. KGN-GC treated with extremely low or sufficient doses of 1,25-(OH)_2_-D3 in the presence of Fsk, were immunoprecipitated with either anti-VDR or anti-RXR antibodies. The VDR-IP pulls down all forms of VDR:homodimers, heterodimers and solo receptor as does the RXR-IP. The immuno-precipitated samples were then immunoblotted using either anti-VDR or anti-RXR antibodies to assess just the heterodimer association between RXR and VDR in the relevant IP samples (figure 6). The subsequent western blot membranes were analysed in a variety of ways (tables 4a and 4b).There was a significant reduction in levels of VDR associated with RXR-IP in the presence of VD [20nM] combined with Fsk compared to basal, FSk alone or Fsk+ 0.02nM 1,25-(OH)_2_-D3 (table 4a) However, there was no change in RXR levels associated with VDR-IP (table 4b).

**Fig 6:**
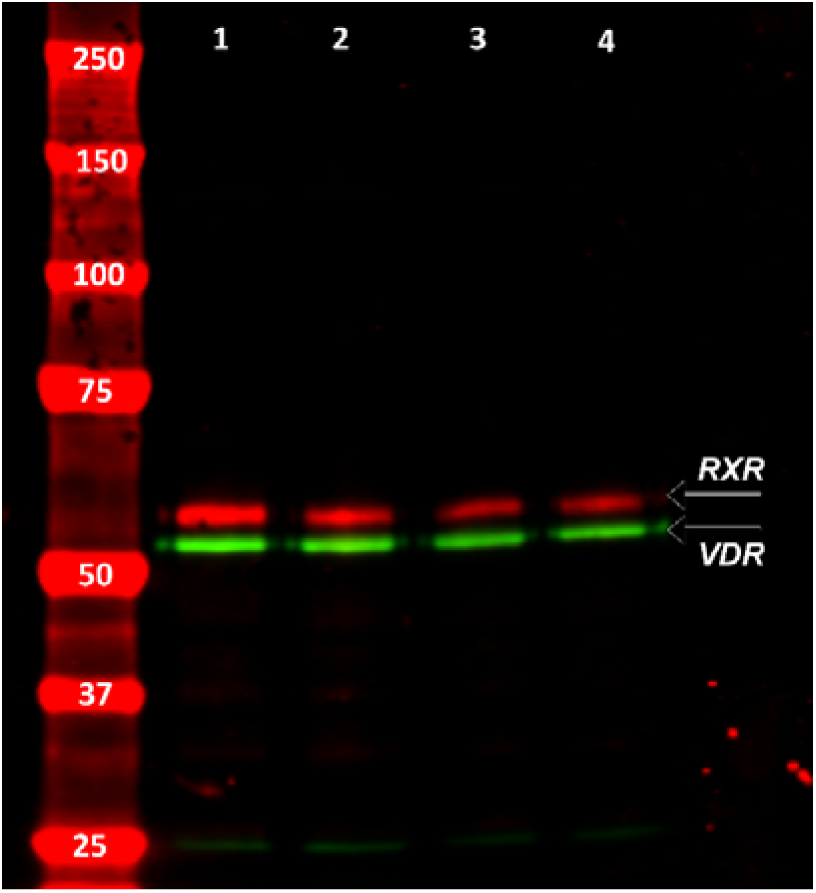
Representative image of western blot of protein from KGN cells cultured ± Fsk and VD at 0.02 and 20nM; immuno-precipitated with anti-VDR antibody (1:1000) and western blotted with anti-RXR antibody (1:1000) to detect RXR:VDR heterodimers. Equal quantities of protein were loaded onto each lane to allow for comparisons. *Lanes: 1=control, 2=Fsk, 3=Fsk+VD20,4=Fsk+VD0.02 Red band=anti-RXRα, Green band=anti-VDR (IP-VDR)*.

## Discussion

We have discovered widespread distribution of VDR in the human ovary, pointing to a likely critical role for VD in female reproduction. We have demonstrated for the first time, a clear relationship between increasing VDR expression in theca and follicle size, highlighting a possible important functional role for Vitamin D in antral follicle progression.

Our findings in human theca were supported by the results of histological analysis of mouse ovarian follicles which showed an increase in the intensity of VDR expression in theca of LAFs. That activation of these receptors is playing a role in follicle growth was demonstrated by the fact that VD3 supplementation promoted the survival and growth of *in vitro* cultured preantral follicles to the antral stage in the primate ovary (Xu J, 2018). Our finding of protein expression of VDR in human cortex and stroma was also matched by extensive and uniform expression of VDR protein in granulosa cells, cortex, and stroma of mouse ovaries, further indicating an important role for VD throughout the ovary. This is also supported by the fact that female VDR knockout mice have impaired folliculogenesis with no progression beyond primary and secondary stages and low oestradiol levels (Yoshizawa T, 1997).

The detection of *CYP27B1* mRNA expression in human ovarian stroma was intriguing as it indicated that the human ovary is capable of being an extra-renal site of active 1,25-(OH)_2_-D3 synthesis, as supported by studies in other species (Xu J *et al*, 2018). The contribution of any locally produced bioactive 1,25-(OH)_2_-D3 is difficult to determine, but could play an important role in counteracting systemic VD deficiency. Active 1,25-(OH)_2_-D3 negatively regulates expression of CYP27B1, hence regulating its own circulating concentrations (Bouillon R *et al*, 2008).

This increasing expression of VDR in the growing follicle at the LAF stage corresponds to increasing steroidogenesis and the acquisition of LH receptors. It had been documented that incubation of human cumulus GC with VD at normal levels affects steroid output (Merhi Z *et al*, 2014), but we wished to investigate the extent of and mechanisms by which exposing cells to a 1,25-(OH)_2_-D3-deficient environment affected steroidogenesis. To that end we incubated cells with 20, 2, 0.2 or 0.02nmol/L (since serum levels <50nmol/L is defined as severe deficiency) (Holick MF *et al*, 2011). A recent study of women of childbearing age in rural northern China found the prevalence of severe VD deficiency was 16% in the 1151 women studied, with a median serum level of only of 5.63ng/ml (equivalent to 14nmol/L) (Lin S *et al*, 2021); indicating that severe hypovitaminosis of this magnitude occurs in the population. Interestingly, 1,25-(OH)_2_-D3 is established to act at nanomolar, or even picomolar, concentrations, as a direct regulator of specific target genes in VDR-expressing cell and tissues. It also binds to VDR with an affinity of 0.1nM (reviewed in Carlsberg, 2022).

In GC, gonadotrophins stimulate oestrogen synthesis via cAMP-mediated signalling, and it is well-established that a cAMP-response element (CRE) sequence has been identified within promoter II (PII); the predominant aromatase promoter used in the ovary (Michael MD, 1997). Changes in the 1,25-(OH)_2_-D3 environment had no effect on E2 production from GC of either SAFs/LAFs, in the presence or absence of FSH and FSH responsiveness was retained. Conversely in <u>luteinised</u> granulosa cells (GLCs), 1,25-(OH)_2_-D3 did prevent the LH-mediated stimulation of E2 production. It is without doubt that 1,25-(OH)_2_-D3 can decrease E2 production as the same effect was observed with the VD analogue in GLCs.

This would imply that attenuation by VD only occurred when GC acquired LHR or else is dependent on levels of cAMP generated, since LH stimulates more cAMP activity than FSH (Aharoni D *et al*, 1995). This is supported by the observation that extremely low doses of 1,25-(OH)_2_-D3 down-regulated forskolin-stimulated aromatase expression and activity in KGN cells (which do not have LHR), an effect that was lost once dosing levels were increased.

This contrasted with theca, where a low 1,25-(OH)_2_-D3 significantly attenuated androstenedione production from theca of LAF (15-22mm) but not of SAF (<10mm). Surprisingly, 1,25-(OH)_2_-D3 had no effect on progesterone or 17-OH-P production from theca of all sized follicles indicating that this inhibitory action presumably occurs in the *CYP17* pathway, but only in LAFs when they have upregulated VDR expression. Like observations in theca, there was no effect of 1,25-(OH)_2_-D3 on progesterone production from GLCs.

The bi-phasic dose-response effects of 1,25-(OH)_2_-D3 on aromatase expression, was a direct influence on the transcriptional activity of PII, as seen in the PII transfection assay results. Analysis of PII has revealed the presence of two VDRE - proximal and distal, with an overlap between the proximal VDRE and CRE (Krishnan AV *et al*, 2010). A key factor could be the mechanism of ligand-binding of VD to its receptor. VDR functions as both a monomer, homodimer, and a heterodimer with RXR, and the alteration of the ratio of these complexes within a cell is dependent on the amount of VD ligand present (Cheskis B *et al*, 1994). Depending on the proportion of each, these complexes will bind to VDRE on genes and attract either co-repressors/activators to enhance or suppress gene expression and activity. Increased levels of VD decreased the amount of DNA-bound VDR homodimer complexes and promote the formation of VDR-RXR heterodimers (Carlsberg C, 2022; Cheskis B *et al*, 1994; Haussler MR *et al*, 2013).

<u>Unliganded</u> VDR-RXR heterodimers are initially bound to a VDRE and recruit co-repressor complexes, which prevents basal transcription through the activity of histone deacetylase. Once sufficient ligand has bound, the repressors are substituted by co-activator complexes, allowing gene transcription to commence (Carlsberg C, 2022; Dwivedi PP *et al*, 1998; Perissi V *et al*, 2010). The results of our IP experiments showed a difference in the levels of VDR associated with immunoprecipitated RXR, at different concentrations of VD ligand in the presence of forskolin. In the presence of higher concentrations of VDR ligand there appears to be less VDR associated with immunoprecipitated RXR. However, it must be pointed out that it was not possible from our experiments to determine the proportion of heterodimers in the cytoplasmic or nuclear compartments (and hence non-genomic/genomic forms) as we used whole cell lysates. To answer this, we would need to carry out further experiments (beyond scope of this study) to explore VDR sub-cellular trafficking between nuclear and cytoplasmic compartments and components of the repressor/activator complexes bound to chromatin. Interestingly, the ability of forskolin to alter the association of the intracellular dimers would indicate the presence of a ligand-independent cAMP activated pathway outside the nucleus (Luk J *et al*, 2012); (Haussler MR *et al*, 2013) which could account for the different outcomes observed when using either FSH or LH.

As stated earlier, women with PCOS commonly present with increased serum levels of AMH (reflecting the pool of stalled SAF) and hyperinsulinemia, both of which are also linked to VD status (Luk J, 2012; Lorenzen M. B., 2017). To investigate this link, we replicated hyperinsulinemia and insulin resistance *in vitro* by chronically exposing KGN cells to insulin at either post-prandial (10ng/ml) or hyperinsulinemic (100ng/ml) doses. The HI dose downregulated expression of insulin receptor mRNA by more than 50% reproducing insulin resistance, whereas there was no significant effect noted at post-prandial treatment levels. To our surprise, culture of cells in an extremely low and deficient VD environment caused an even further attenuation of insulin receptor expression in the presence of both doses of insulin. This effect only occurred in the presence of insulin, as 1,25-(OH)_2_-D3 alone had no effect on insulin receptor expression. Systemic VD deficiency is clearly linked to a reduction in insulin sensitivity (independent of BMI) as others have shown (Muscogiuri G *et al*, 2017), and this insulin receptor reduction may be a contributory mechanism. Interestingly, in the same cells a combination of insulin and 1,25-(OH)_2_-D3 had no effect on aromatase, unlike in the forskolin experiments, again indicating that this was a cAMP-driven process, possibly linked to overlap between VDRE and CRE on PII as described previously.

It is well established that follicle progression in the normal human ovary requires down-regulation of AMH expression to permit FSH-driven activity (Pellatt L *et al*, 2010). The IHC analysis of mouse ovaries also revealed high AMH expression in SAF which was substantially reduced in pre-ovulatory follicles. This is thought to normally occur via LH down-regulating AMH expression (Pierre A *et al*, 2013). Using forskolin instead of LH to achieve down-regulation, we demonstrated that this reduction in AMH was potentiated by 1,25-(OH)_2_-D3 [0.02nM] with a further reduction occurring with higher doses of 1,25-(OH)_2_-D3 [20nM]. Interestingly treatment with 1,25-(OH)_2_-D3 alone had no significant effect on AMH mRNA levels, indicating again that this only occurs when the cAMP pathway is activated.

In hen GC from 3-5mm and 6-8mm follicles, vitamin D3 was shown to substantially decrease AMH mRNA expression, with a more robust effect seen in the LAF (Wojtusik J, 2012). In addition, Mehri *et al* showed that in AF (<14mm) from women with insufficient/deficient follicular fluid levels of 25-OH-D3, there was significantly higher AMHR-II mRNA level compared to those with normal VD levels (Merhi Z *et al*, 2014). In the same study, 25-OH-D3 was shown to decrease AMH-mediated pSMAD 1/5/8 nuclear localisation in cumulus GC, indicative of reduced AMH signalling. Hence at a local ovarian level, our data and other studies, clearly showed that VD is involved in AMH regulation and expression, which could impact on follicle progression particularly in conditions such as PCOS. At a systemic level however, evidence for correlations between serum vitamin D and serum AMH levels are contradictory and dependent on a woman’s ovulatory status (Moridi I *et al*, 2020).

To summarise, we have shown that Vitamin D clearly has a role to play in the theca and granulosa cell function and hence growth of ovarian follicles, as shown by the increased expression of VDR in follicles of increasing size. In addition, we have shown that Vitamin D may promote follicle progression by downregulating the expression of AMH, thereby reducing AMH’s well-documented inhibitory effect on follicle growth (Pellatt L *et al*, 2010). Vitamin D is known to affect steroidogenesis and we have demonstrated that levels of 1,25-(OH)_2_-D3 equivalent to hypovitaminosis, inhibited thecal production of androstenedione. In addition, extremely low levels of vitamin D had an attenuating effect on cAMP-driven aromatase expression, which translated to decreased E2 production. Encouragingly this reduction in E2 is reversed as levels of 1,25-(OH)_2_-D3 increased; apart from in the presence of LH in luteinised GC, which could have consequences for regular ovulation in women with severe Vitamin D deficiency. For the first time we have demonstrated that deficient levels of 1,25-(OH)_2_-D3 also down-regulated insulin receptor expression, potentially reducing insulin sensitivity. This could have serious implications for women with hyperinsulinemia and insulin resistance, typically seen in PCOS; indicating that insulin resistant women should try and maintain sufficient levels of systemic vitamin D for regular ovarian function. The ability of the ovary to make local bioactive 1,25-(OH)_2_-D3, was demonstrated by the expression of *CYP27B1.* This together with the upregulation of VDR expression in all ovarian cellular compartments, could potentially counteract the effect of systemic VD deficiency and protect the local ovarian environment. To conclude a severely deficient VD environment (<2nM or <1ng/ml) could contribute to impaired ovarian cell function and hence potentially affect folliculogenesis/ovulation, but levels associated with mild deficiency may have less impact, apart from in the presence of hyperinsulinemia and insulin resistance.

## Statement regarding use of archived tissue

Research and the Human Tissue Act 2004 - Consent IS REQUIRED to use and store tissues for research; UNLESS: The relevant material is classed as an existing holding i.e., held prior to 1st September 2006 and/or the relevant material is imported.

## Acknowledgements

We would like to thank the doctors and nurses at Malta Medical School, Msida who provided us with the ovaries and laboratory space; in particular Dr Ray Galea and Prof. Mark Brincat. Additionally, we would like to thank Mr Michael Lacey for his technical assistance with various aspects of the project; along with Ms Yvette Bland for her help with the IHC of mouse ovaries. Particular thanks must go to Erin Vang who helped with the retrieval and conversion of Statview Data files to Excel. We also acknowledge the experimental contribution of intercalated and BSc students at St. George’s University of London – Angus Mansfield, Andrew Standing and Ella Jameson.

Finally, and most importantly, we would like to acknowledge and thank all the patients who donated tissue and cells which allowed us to conduct this study.

## Supplemental Figure Legends

**Fig S1:** 1,25(OH)_2_D_3_ had no effect on progesterone production from <u>theca</u> taken from LAF (15-22mm, grey bars, *n=5 subjects, total follicles=5*) from normal ovaries. *(ANOVA n/s)*.

**Fig S2:** 1,25(OH)_2_D_3_ had no effect on 17-hydroxy-progesterone (17-OH-P) production from <u>theca</u> taken from LAF (15-22mm, grey bars, *n=5 subjects, total follicles=5*) from normal ovaries. *(ANOVA n/s)*.

**Fig S3:** 1,25(OH)_2_D_3_ had no effect on progesterone production from granulosa-lutein cells *(n=4; ANOVA n/s)*.

## Notes

### Competing Interest Statement

The authors have declared no competing interest.

### Summary of Updates

Change the emphasis from PCOS to mechanisms of VitD's action in ovarian cells. Remove PCOS data for which there was not enough patient samples to assess statistical significance. Explain the methodologies applied in a clearer manner.

## References

Aharoni D, Dantes A, Oren M, Amsterdam. “cAMP-Mediated Signals as Determinants for Apoptosis in Primary Granulosa Cells” Experimental Cell Research (1995) Vol 218 (1):71–282,

Agic A, Xu H, Altgassen C, Noack F, Wolfler MM, Diedrich K, Friedrich M, Taylor RN, Hornung D. “Relative expression of 1,25-dihydroxyvitamin D3 receptor, vitamin D 1 alpha-hydroxylase, vitamin D 24-hydroxylase, and vitamin D 25-hydroxylase in endometriosis and gynecologic cancers. .” Reprod Sci. (2007) 14, (5): 486–97.

Aranow, C.”Vitamin D and the immune system” J.Investig.Med, (2011) 59(6): 881–6

Bouillon R, Carmeliet G, Verlinden L, van Etten E, Verstuyf A, Luderer HF, Lieben L, Mathieu C, Demay M. “Vitamin D and human health: lessons from vitamin D receptor null mice” Endoc. Rev. (2008) 29: 726–776.

Carlsberg, C. (2022). Vitamin D and Its Target Genes. Nutrients, 14(7), 1354.

Cheskis B, Freedman LP. “Ligand modulates the conversion of DNA-bound vitamin D3 receptor (VDR) homodimers into VDR-retinoid X receptor heterodimers.” Mol Cell Biol. (1994) 14, no. 5: 3329–38.

Dilaver N, Pellatt L, Jameson E, Ogunjimi M, Bano G, Homburg R, D Mason H, Rice S. “The regulation and signalling of anti-Müllerian hormone in human granulosa cells: relevance to polycystic ovary syndrome” Hum Reprod. (2019) 34(12): 2467–2479.

Dusso, AS, Brown AJ, Slatopolsky E. “Vitamin D” Am.J.Physiol. (2005) 289(1):F8–28

Dwivedi PP, Muscat GE, Bailey PJ, Omdahl JL, May BK. “Repression of basal transcription by vitamin D receptor: evidence for interaction of unliganded vitamin D receptor with two receptor interaction domains in RIP13delta1.” J Mol Endocrinol. (1998) 20, no. 3: 327–35.

Eftekhar, M., Mirhashemi, E. S., Molaei, B., & Pourmasumi, S. “Is there any association between vitamin D levels and polycystic ovary syndrome (PCOS) phenotypes?” Archives of endocrinology and metabolism 64, no. 1 (2020): 11–16.

Gilling-Smith C, Willis DS, Beard RW, Franks S. “Hypersecretion of androstenedione by isolated thecal cells from polycystic ovaries.” Journal of Clinical Endocrinology and Metabolism 79 (1994): 1158–1165.

Grzesiak M, Knapczyk-Stwora K, Slomczynska M. Vitamin D3 in ovarian antral follicles of mature gilts: Expression of its receptors and metabolic enzymes, concentration in follicular fluid and effect on steroid secretion in vitro. Theriogenology. 2021;160:151–160.

Haussler MR, Whitfield GK, Kaneko I, Haussler CA, Hsieh D, Hsieh JC, Jurutka PW. “Molecular mechanisms of vitamin D action”. Calcif Tissue Int. 2013 Feb;92(2):77–98

Hillier S G, Van de Boogaard A M, Reichert L E Jr, Van Hall E V. “Alterations in granulosa cell aromatase activity accompanying preovulatory follicular development in the rat ovary with evidence that 5 apha-reduced C19 steroids inhibit the aromatase.” J Endocrinol 84 (1980): 409–19.

Holick, MF. “Vitamin D deficiency” NEJM (2007)357(3): 266–281.

Holick MF, Binkley NC, Bischoff-Ferrari HA, Gordon CM, Hanley DA, Heaney RP, Murad MH, Weaver CM, Endocrine Society. “Evaluation, treatment, and prevention of vitamin D deficiency: an Endocrine Society clinical practice guideline.” J Clin Endocrinol Metab. 96, no. 7 (2011): 1911–30.

Holick MF, Binkley NC, Bischoff-Ferrari HA, Gordon CM, Hanley DA, Heaney RP, M. Murad H, Weaver CM. “Guidelines for Preventing and Treating Vitamin D Deficiency and Insufficiency Revisited.” The Journal of Clinical Endocrinology & Metabolism 97, no. 4 (2012): 1153–1158.

Krishnan AV, Moreno J, Nonn L, Malloy P, Swami S, Peng L, Peehl DM, Feldman D. “Novel pathways that contribute to the anti-proliferative and chemopreventive activities of calcitriol in prostate cancer.” J Steroid Biochem Mol Biol 103, no. (3-5) (2007): 694–702.

Krishnan AV, Swami S, Peng L, Wang J, Moreno J, Feldman D. “Tissue-selective regulation of aromatase expression by calcitriol: implications for breast cancer therapy. .” Endocrinology 151, no. 1 (2010): 32–42.

Krul-Poel YHM, Koenders PP, Steegers-Theunissen RP, Ten Boekel E, Wee MMT, Louwers Y, Lips P, Laven JSE, Simsek S. “Vitamin D and metabolic disturbances in polycystic ovary syndrome (PCOS): A cross-sectional study. .” PLoS One 13, no. 12 (2018): e0204748.

Łagowska, K., Bajerska, J., & Jamka, M. “The Role of Vitamin D Oral Supplementation in Insulin Resistance in Women with Polycystic Ovary Syndrome: A Systematic Review and Meta-Analysis of Randomized Controlled Trials.” Nutrients, 10, no. 11 (2018): 1637.

Lerchbaum, E, and B Obermayer-Pietsch. “Vitamin D and fertility: a systematic review” EJE (2012) 166(5):765–78.

Lin S, Jiang L, Zhang Y, Chai J, Li J, Song X, Pei L. “Socioeconomic status and vitamin D deficiency among women of childbearing age: a population-based, case-control study in rural northern China. .” BMJ Open. 11, no. 3 (2021): e042227.

Lorenzen, M., Boisen, I. M., Mortensen, L. J., Lanske, B., Juul, A., & Blomberg Jensen, M. “Reproductive endocrinology of vitamin D.” Molecular and cellular endocrinology 453 (2017): 103–112.

Luderer HF, Gori F, Demay MB. “Lymphoid enhancer-binding factor-1 (LEF1) interacts with the DNA-binding domain of the vitamin D receptor” J Biol Chem. (2011) 286(21):18444–51.

Luk J, Torrealday S, Neal Perry G, Pal L. “Relevance of vitamin D in reproduction. .” Hum Reprod. 27, no. 10 (2012): 3015–3027.

Maestro B, Dávila N, Carranza MC, Calle C. “Identification of a Vitamin D response element in the human insulin receptor gene promoter.” J Steroid Biochem Mol Biol., 2003: 223–30.

Mason, H. D., Martikainen, H., Beard, R. W., Anyaoku, V., & Franks, S. “Direct gonadotrophic effect of growth hormone on oestradiol production by human granulosa cells in vitro.” The Journal of endocrinology 126, no. 3 (1990): R1–R4.

Mason, H. D., Cwyfan-Hughes, S. C., Heinrich, G., Franks, S., & Holly, J. M. “Insulin-like growth factor (IGF) I and II, IGF-binding proteins, and IGF-binding protein proteases are produced by theca and stroma of normal and polycystic human ovaries. .” The Journal of clinical endocrinology and metabolism 81, no. 1 (1996): 276–284.

McCaig, C., Perks, C. M., & Holly, J. M. “Signalling pathways involved in the direct effects of IGFBP-5 on breast epithelial cell attachment and survival.” Journal of cellular biochemistry 84, no. 4 (2002): 784–794.

Menichini D., Facchinetti F. “Effects of vitamin D supplementation in women with polycystic ovary syndrome: A review.” Gynecol. Endocrinol. 36 (2020): 1–5.

Merhi Z, Doswell A, Krebs K, Cipolla M. “Vitamin D alters genes involved in follicular development and steroidogenesis in human cumulus granulosa cells.” J Clin Endocrinol Metab. 99, no. 6 (2014): E1137–45.

Michael MD, Michael LF, Simpson ER. “A CRE-like sequence that binds CREB and contributes to cAMP-dependent regulation of the proximal promoter of the human aromatase P450 (CYP19) gene.” Mol Cell Endocrinol. 134, no. 2 (1997): 147–56.

Moridi I, Chen A, Tal O, Tal R. “The Association between Vitamin D and Anti-Müllerian Hormone: A Systematic Review and Meta-Analysis.” Nutrients. 12, no. 6 (2020): 1567.

Muscogiuri G, Altieri B, de Angelis C, Palomba S, Pivonello R, Colao A, Orio F. “Shedding new light on female fertility: The role of vitamin D.” Rev Endocr Metab Disord. 18, no. 3 (2017): 273–283.

Nestler J.E., Jakubowicz D.J., de Vargas A.F., Brik C., Quintero N., Medina F. “Insulin stimulates testosterone biosynthesis by human thecal cells from women with polycystic ovary syndrome by activating its own receptor and using inositolglycan mediators as the signal transduction system.” J. Clin. Endocrinol. Metab. 83 (1998): 2001–2005.

Nishi Y, Yanase T, Mu Y, Oba K, Ichino I, Saito M, Nomura M, Mukasa C, Okabe T, Goto K, Takayanagi R, Kashimura Y, Haji M, Nawata H. “Establishment and characterization of a steroidogenic human granulosa-like tumor cell line, KGN, that expresses functional follicle-stimulating hormone receptor.” Endocrinology. 2001 Jan;142(1):437–45.

Pellatt L, Rice S, Mason HD. “Anti-Müllerian hormone and polycystic ovary syndrome: a mountain too high? .” Reproduction 139, no. 5 (2010): 825–33.

Perissi V, Jepsen K, Glass CK, Rosenfeld MG. “Deconstructing repression: evolving models of co-repressor action.” Nat Rev Genet. 11, no. 2 (2010): 109–23.

Pierre A, Peigné M, Grynberg M, Arouche N, Taieb J, Hesters L, Gonzalès J, Picard JY, Dewailly D, Fanchin R, Catteau-Jonard S, di Clemente N. “Loss of LH-induced down-regulation of anti-Müllerian hormone receptor expression may contribute to anovulation in women with polycystic ovary syndrome. .” Hum Reprod. 28, no. 3 (2013): 762–9.

Pilz S, Zittermann A, Trummer C, et al. “Vitamin D testing and treatment: a narrative review of current evidence.” Endocr Connect. 8, no. 2 (2019): R27–R43.

Rice S, Mason HD, Whitehead SA. “Phytoestrogens and their low dose combinations inhibit mRNA expression and activity of aromatase in human granulosa-luteal cells.” J Steroid Biochem Mol Biol. 101, no. 4-5 (2006): 216–25.

Rice, S., Elia, A., Jawad, Z., Pellatt, L., & Mason, H. D. “Metformin inhibits follicle-stimulating hormone (FSH) action in human granulosa cells: relevance to polycystic ovary syndrome.” The Journal of clinical endocrinology and metabolism 98, no. 9 (2013): E1491–E1500.

Schuster I. “Cytochromes P450 are essential players in the vitamin D signaling system.” Biochim Biophys Acta. 1814, no. 1 (2011): 186–199.

Tiejun Sun, Ying Zhao, David J. Mangelsdorf, Evan R. Simpson. “Characterization of a Region Upstream of Exon I.1 of the Human CYP19 (Aromatase) Gene That Mediates Regulation by Retinoids in Human Choriocarcinoma Cells.” Endocrinology, 1998: 1684–1691.

Wang L, Lv S, Li F, Yu X, Bai E, Yang X. “Vitamin D Deficiency Is Associated With Metabolic Risk Factors in Women With Polycystic Ovary Syndrome: A Cross-Sectional Study in Shaanxi China. .” Front Endocrinol (Lausanne) 11 (2020): 171.

Willis DS, Mason HD, Gilling-Smith C, Franks S. “Modulation by insulin of follicle stimulating hormone and luteinising hormone actions in human granulosa cells of normal and polycystic ovaries.” J Clin Endo Metab 81 (1996): 302–309.

Wojtusik J, Johnson PA. “Vitamin D regulates anti-Mullerian hormone expression in granulosa cells of the hen. .” Biol Reprod. 86, no. 3 (2012): 91.

Wright, R. J., Holly, J. M., Galea, R., Brincat, M., & Mason, H. D. “Insulin-like growth factor (IGF)-independent effects of IGF binding protein-4 on human granulosa cell steroidogenesis.” Biology of reproduction, 67, no. 3 (2002): 776–781.

Xu J, Hennebold JD, Seifer DB. Direct vitamin D3 actions on rhesus macaque follicles in three-dimensional culture: assessment of follicle survival, growth, steroid, and antimüllerian hormone production. Fertil Steril. 2016;106(7):1815–1820.e1.

Xu J, Lawson MS, Xu F, Du Y, Tkachenko OY, Bishop CV, Pejovic-Nezhat L, Seifer DB, Hennebold JD. “Vitamin D3 Regulates Follicular Development and Intrafollicular Vitamin D Biosynthesis and Signaling in the Primate Ovary. .” Front Physiol. 14, no. 9 (2018): 1600.

Yoshizawa T, Handa Y, Uematsu Y, Takeda S, Sekine K, Yoshihara Y, Kawakami T, Arioka K, Sato H, Uchiyama Y, Masushige S, Fukamizu A, Matsumoto T, Kato S. “Mice lacking the vitamin D receptor exhibit impaired bone formation, uterine hypoplasia and growth retardation after weaning.” Nat Genet. 16, no. 4 (1997): 391–6.

Zella LA, Kim S, Shevde NK, Pike JW. “Enhancers located in the vitamin D receptor gene mediate transcriptional autoregulation by 1,25-dihydroxyvitamin D3.” J Steroid Biochem Mol Biol. 103, no. 3-5 (2007): 435–9.

